# Bridges instead of boats? The Mla system of diderm Firmicute *Veillonella parvula* reveals an ancestral transenvelope core of phospholipid trafficking

**DOI:** 10.1101/2023.06.30.547184

**Authors:** Kyrie P. Grasekamp, Basile Beaud, Najwa Taib, Bianca Audrain, Benjamin Bardiaux, Yannick Rossez, Nadia Izadi-Pruneyre, Maylis Lejeune, Xavier Trivelli, Zina Chouit, Yann Guerardel, Jean-Marc Ghigo, Simonetta Gribaldo, Christophe Beloin

## Abstract

Despite extensive characterisation of envelope biogenesis systems in diderm bacteria, glycerophospholipid (GPL) trafficking remains poorly understood, and has only been studied in a handful of model species. Within the Proteobacteria, the maintenance of lipid asymmetry (Mla) system facilitates retrograde GPL trafficking via six proteins, MlaA-F. GPLs are extracted from the outer leaflet of the outer membrane by the lipoprotein MlaA which associates with porin trimers, then shipped through the periplasmic space by the chaperone MlaC, which finally delivers GPLs to the inner membrane complex formed by MlaBDEF. Here, we investigate GPL trafficking in *Veillonella parvula*, a diderm member of the Firmicutes which encodes an Mla system devoid of MlaA and MlaC. *V. parvula* Δ*mla* mutants display phenotypes characteristic of disrupted lipid asymmetry such as hypervesiculation and detergent hypersensitivity, and lipid content analysis from outer membrane vesicles reveals an enrichment for the major lipid component phosphatidylethanolamine. Interestingly, suppressor analysis identifies mutations in *tamB* that rescue detergent hypersensitivity and hypervesiculation of Δ*mla* strains, supporting the involvement of these two systems in antagonistic GPL trafficking functions across diverse bacterial lineages. A combination of structural modeling and subcellular localisation assays shows that MlaD_Vp_ is longer than in classical diderm models and forms a transenvelope bridge, encoding both an inner membrane-localised MCE domain and an outer membrane ß-barrel. These results strongly suggest that *V. parvula* possesses a minimal Mla system for GPL trafficking, replacing the need for chaperones and outer membrane lipoproteins by directly connecting the two membranes. Finally, phylogenomic analysis indicates that this MlaEFD self-contained architecture is widely distributed in diderm bacteria and most likely represents the ancestral functional core of the Mla system, which subsequently increased in complexity in Proteobacteria and closely related phyla following the emergence of MlaABC. Our work broadens the diversity of current models of GPL trafficking in diderm bacteria, challenging the paradigm set by classical models and shedding light on the evolution of a crucial system in the biogenesis and maintenance of the bacterial outer membrane.

## Introduction

The diderm bacterial envelope is a complex structure consisting of two membranes: a cytoplasmic -or inner-membrane (IM) composed primarily of glycerophospholipids (GPLs), and an outer membrane (OM) which exhibits lipid asymmetry, comprising GPLs in its inner leaflet and primarily lipopolysaccharide (LPS) molecules in its outer leaflet. This asymmetry manifests as a robust exclusion barrier against a range of antibiotics and host components, such as vancomycin and bile salts^1, 2^. Although GPLs are one of the most ubiquitous amphipathic components of the diderm envelope, their transport systems remain poorly understood, and most current information stems from a handful of model bacterial species. Recent studies implicate large AsmA-like proteins, such as TamB and YhdP, in the mediation of anterograde GPL trafficking^3–5^, whilst MCE proteins – namely MlaD within the maintenance of lipid asymmetry (Mla) machinery - are proposed to facilitate retrograde GPL trafficking^6, 7^. In *E. coli,* the Mla system is composed of six proteins **(Fig 1)**: MlaBDEF form the IM complex^8, 9^, MlaC is the periplasmic chaperone^6, 10^, and MlaA is the OM lipoprotein that associates with porin trimers^11^. Despite extensive research of the Mla system within model diderms for over a decade, several mechanistic details – including substrate specificity and even directionality, until recently - remain controversial^9, 12–14^.

**Figure 1:**
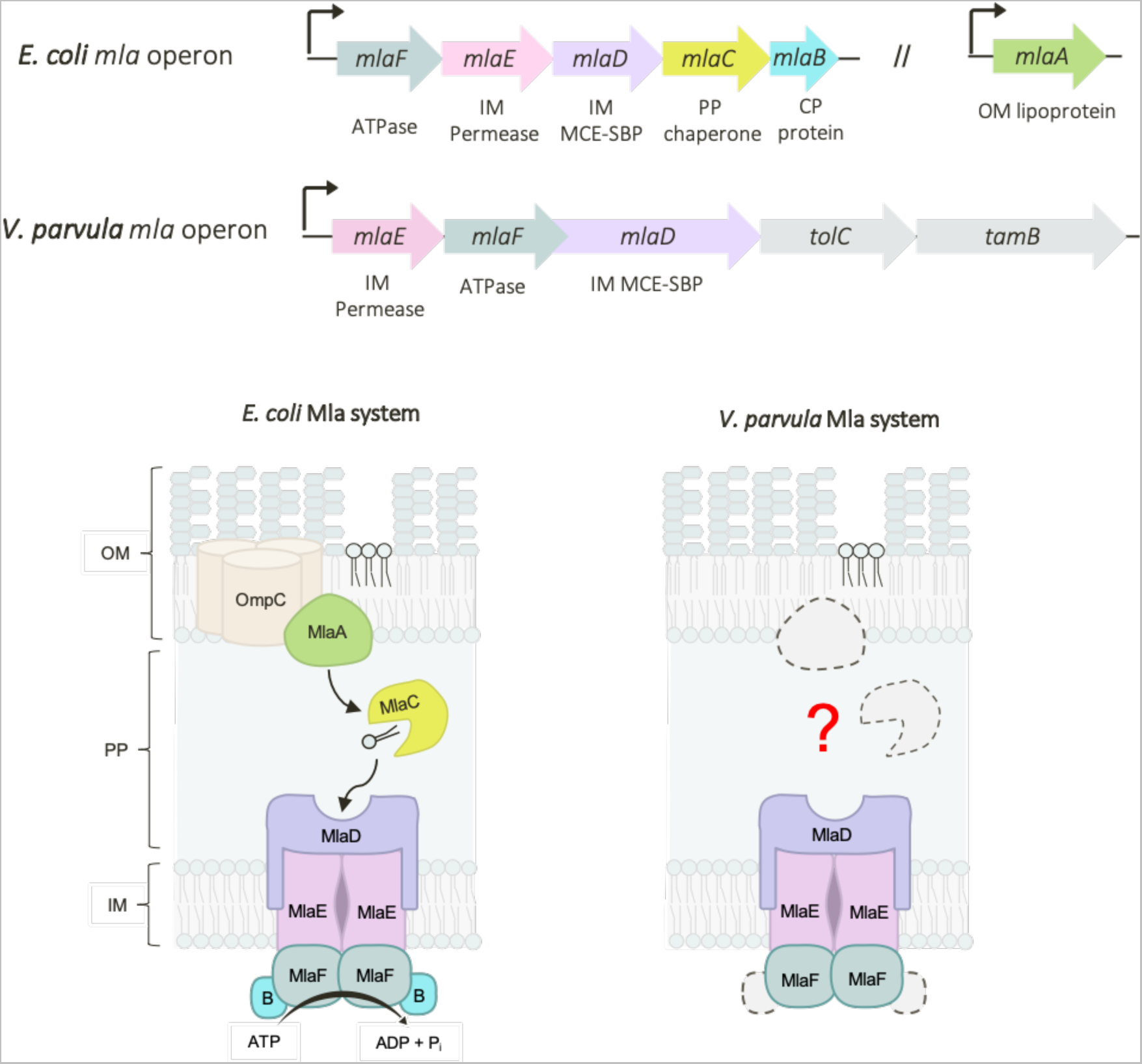
Comparison of Mla operons in *V. parvula* and *E. coli*. *In E. coli*, the Mla system is encoded by *mlaB-F*, present in a single operon, and *mlaA* at a separate locus. MlaBDEF form the IM complex, an ABC transporter in which MlaD is the MCE-domain substrate binding protein (SBP); MlaC is the periplasmic chaperone, and MlaA is the OM lipoprotein that associates with OmpC trimers. In *V. parvula*, only homologues of *mlaDEF* were identified. These genes are encoded together in an operon, immediately upstream of a homologue of *tolC* and *tamB*. OM= outer membrane; PP = periplasm; IM = inner membrane; CP = cytoplasm.

**Figure 2:**
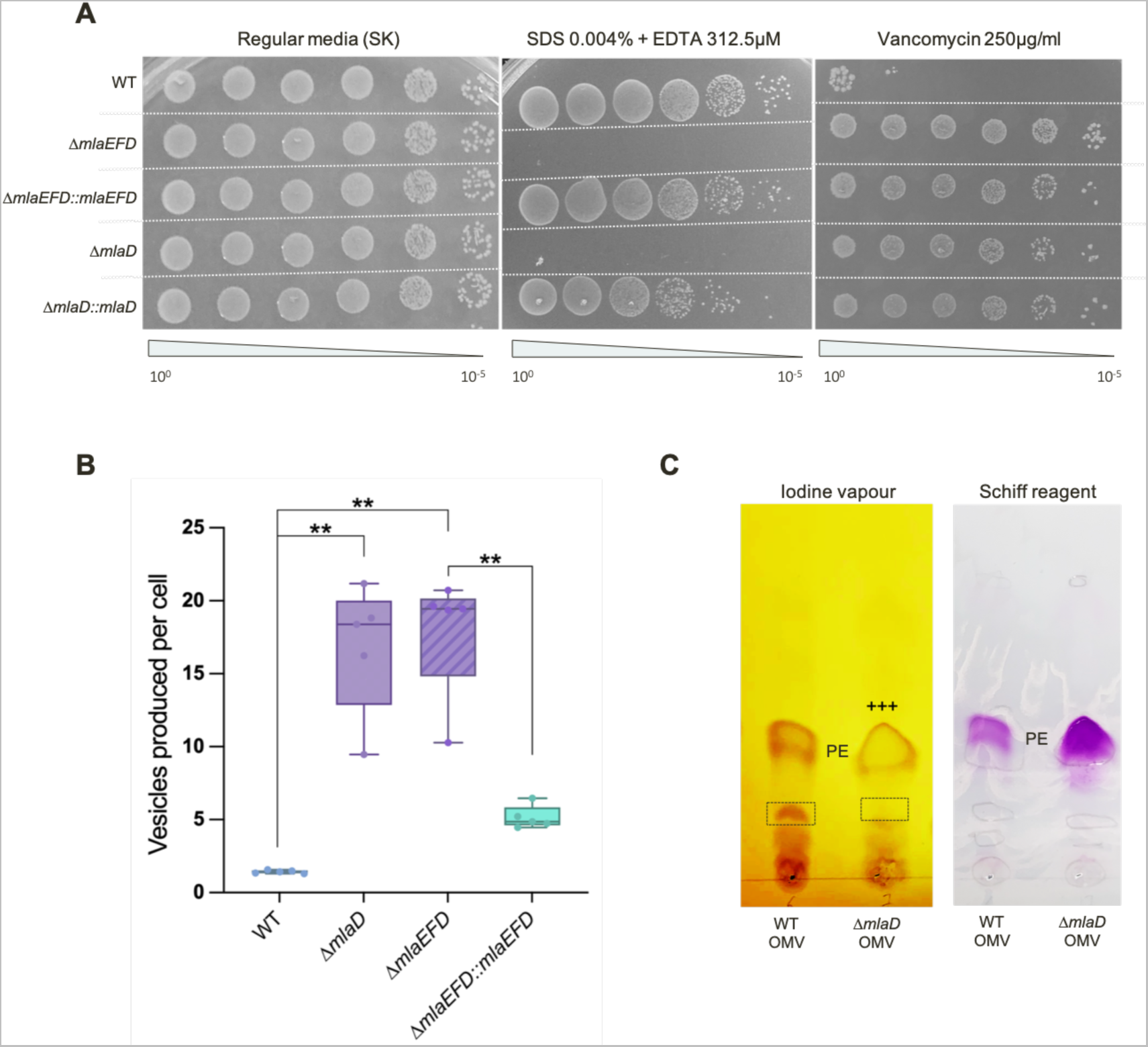
Phenotypic characterisation of Δ*mla* mutants reveals detergent hypersensitivity and hypervesiculation. **A) Efficiency of plating assay shows SDS / EDTA sensitivity of Δ*mla* deletion strains**. Overnight cultures were adjusted for OD_600_ and serial diluted onto SK media supplemented with SDS / EDTA and vancomycin. Δ*mla* mutants display hypersensitivity to SDS / EDTA, and a strong resistance to vancomycin. Complementation of the detergent hypersensitivity was performed with an aTc-inducible vector pRPF185. **B**) **Hypervesiculation of Δ*mla* deletion strains.** Outer membrane vesicles (OMVs) produced by the WT, Δ*mlaD,* Δ*mlaEFD* and complemented strains were quantified via NanoFCM (see methods) and used to generate a ratio of hypervesiculation. The WT produces a similar number of vesicles to cells, resulting in a ∼1:1 ratio, whilst the Δ*mla* deletion strains produce ∼17-fold more vesicles than the WT. This phenotype can be significantly rescued, reducing vesicle production to ∼5-fold in the Δ*mlaEFD::mlaEFD* strain. 5 biological replicates and 3 technical replicates were tested per strain; n = 5; significance calculated by Mann Whitney U, p = 0.0079; two-tailed p-value. **C) Enrichment of phosphatidylethanolamine (PE) in OMVs produced by Δ*mlaD*.** Thin layer chromatography (TLC) of lipid extracts from OMVs produced by both the WT and Δ*mlaD* strains show a ∼40% (+/- 16%) enrichment for PE in the vesicles produced by the Δ*mla* mutants when stained with iodine vapour (labelled +++, quantified by ImageJ over 3 biological replicates). We also notice a decrease in the abundance of another as yet unidentified lipid species, highlighted in the black box. Schiff reagent staining shows a ∼130% increase in the abundance of plasmenyl PE in Δ*mlaD* OMV lipid extracts.

Here, we investigate GPL trafficking in *Veillonella parvula,* a genetically tractable member of the Firmicutes (Negativicutes) which presents a diderm envelope with LPS^15, 16^. *V. parvula* is phylogenetically distant from classical models such as *E. coli* as it belongs to the Terrabacteria, one of the two large clades into which Bacteria are divided (containing phyla such as Actinobacteria, Cyanobacteria and Candidate Phyla Radiation (CPR)), and whose cell envelopes are largely understudied^16, 17^. Remarkably, the Firmicutes is the only phylum so far identified to contain a mixture of both monoderm and diderm clades, offering an ideal platform on which to study both the monoderm-diderm transition^18^ and the diversity of OM biogenesis systems^19^. We previously demonstrated that this diderm architecture is an ancestral feature, present in both the ancestor of the Firmicutes and the Last Bacterial Common Ancestor (LBCA)^15, 17–19^, an inference also supported by a recent study^20^. Studying diderm Firmicutes can therefore shed light on the evolution of ancestral systems in the biogenesis and maintenance of the bacterial OM. Interestingly, the *V. parvula* genome contains an operon embedded in a large OM biogenesis and maintenance cluster^15, 18^ with only three homologues of the Mla system: the IM proteins MlaDEF, directly followed by a homologue of the OM efflux protein TolC^21^, and TamB **(Fig 1)**. This arrangement is conserved in all Negativicutes **(Extended Fig 1)**. However, no homologues of the periplasmic chaperone MlaC or the OM lipoprotein MlaA were identifiable at the sequence level, consistent with the fact that no OM lipoproteins or homologues of the Lol system have so far been identified in *V. parvula*^16, 22^. This raises the question of how these *V. parvula* Mla homologues are involved in retrograde lipid trafficking, and how they can accomplish this function in the absence of MlaABC.

We combine phenotypic characterisation, lipid content analysis, structural modeling and evolutionary reconstructions to show that the three-component Mla system of *V. parvula* is involved in GPL trafficking, and is strikingly different to what has been described so far in model diderms; MlaDEF appear to form a functional ‘minimal’ system, in which a transenvelope MlaD abrogates the requirement for the missing homologues MlaA and MlaC by directly bridging the IM and OM. Further, we show that MlaDEF represent an ancestral core for GPL trafficking that is widely distributed across Bacteria and likely dates back to the LBCA, whereas the presence of MlaABC are an exception, emerging later in the Proteobacteria and other closely related phyla. Finally, and most strikingly, we show that the majority of MlaD sequences across the bacterial kingdom are ‘long’, revealing that the short MlaD employed by Proteobacteria is atypical. Together, our results uncover novel functional information about GPL trafficking in a non-model organism, shedding light on the evolution of the Mla system and challenging the dogma that OM biogenesis systems studied in *E. coli* are representative of all diderms.

## Results

### A "minimal" Mla system in *V. parvula* is involved in GPL trafficking

During a screen to identify genes required for the generation and maintenance of the OM barrier in *V. parvula* SKV38 (see methods), we identified a transposon insertion in a homologue of *mlaD*, and a mixed clone containing an insertion in both *mlaE* and a gene of unknown function, suggesting a role for these *mla* genes in envelope biogenesis. To further investigate the function of the *V. parvula* MlaDEF system, we generated the corresponding single and triple *mlaDEF* deletion mutants and assessed their phenotypes.

Consistent with Mla-defective phenotypes of model diderms such as *E. coli* and *A. baumannii*^7, 12^, all *V. parvula* Δ*mla* strains — with the exception of the Δ*mlaE* mutant that could not be obtained — display hypersensitivity to SDS/EDTA and increased resistance to vancomycin **(Fig 2A)**, with otherwise no differences in viability, morphology, biofilm formation or aggregation **(Extended Fig 2)**. Phenotypes were not additive across single and triple mutants, suggesting these three *mlaDEF* genes work together as part of a single system, further implied by the overlapping start and stop codons of *mlaF* and *mlaD.* SDS/EDTA sensitivity could be complemented in both single and triple mutants, and appeared slightly fuller for the triple mutant, suggesting that correct stoichiometry in the expression of all three genes is important for phenotypic rescue. Δ*mlaEFD* and Δ*mlaD* strains also displayed a ∼17-fold increase in outer membrane vesicle (OMV) production as compared to the WT **(Fig 2B)**, consistent with previously described hypervesiculation phenotypes of Δ*mla* strains in *Vibrio cholerae, Haemophilus influenzae, Bordetella pertussis* and *Neisseria gonorrhoeae*^23–25^, which could be significantly rescued via complementation to a ∼4-fold increase in OMV production as compared to the WT strain. Together, these phenotypes are indicative of a loss of lipid asymmetry^23, 26^. To determine if these changes in membrane permeability and stability are due to a change in quantity or composition of LPS, silver-staining was used to assess relative LPS profiles of these Δ*mla* mutants, revealing no observable differences **(Extended Fig 2B).**

**Figure 3:**
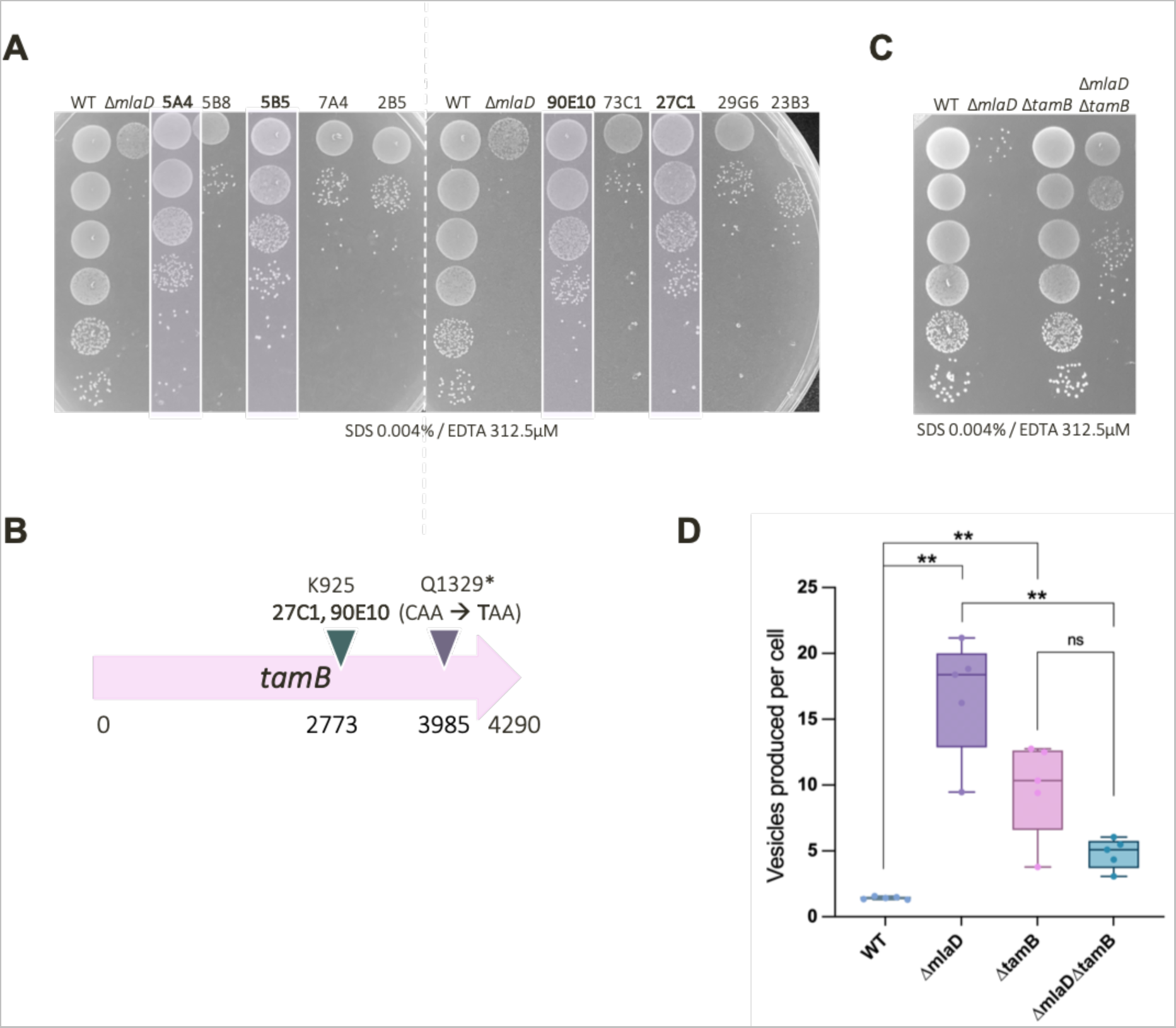
TamB is a suppressor of the Δ*mlaD* OM permeability and hypervesiculation phenotype. **A) Extent of phenotypic rescue of Δ*mlaD* across 10 Tn mutants**. Efficiency of plating assays on SDS 0.004% + EDTA 312.5µM were used to determine the extent of phenotypic rescue by Tn insertions. Only 4 of the 10 Tn insertions could significantly rescue the detergent sensitivity of ΔmlaD: 5A4, 5B5, 27C1 and 90E10. **B) Suppressor mutations identified in *tamB*.** From both spontaneous suppressor analysis, and random Tn insertion suppression analysis, we identified mutations in a homologue of *tamB*. The two identical Tn insertions were at K925, whilst the SNP in the spontaneous suppressor was mapped to Q1329*, generating a STOP codon (CAA → **T**AA). **C) OM permeability of WT, Δ*tamB* and Δ*mla* mutants**. Clean deletion mutants were generated for Δ*tamB, ΔmlaD* and *ΔmlaDΔtamB,* then overnight cultures of these strains and the WT were serially diluted onto SDS / EDTA to assess OM permeability. Δ*tamB* does not display a hypersensitivity to SDS 0.004% / EDTA 312.5µM, whilst Δ*mlaD* is almost unable to grow at this concentration. Deletion of *tamB* in this Δ*mlaD* background partially rescues the detergent hypersensitivity of Δ*mlaD*. **D) Outer Membrane Vesicle (OMV) production of WT, Δ*tamB* and Δ*mla* mutants.** OMV production of WT, Δ*tamB*, Δ*mlaD* and Δ*mlaDΔtamB* strains were analysed via NanoFCM quantification. At least 5 biological replicates and 2-3 technical replicates were tested per strain. Whilst Δ*mlaD* and Δ*tamB* display hypervesiculation, the double mutant (Δ*mlaD*Δ*tamB*) shows a significant reduction in OMV production as compared to Δ*mlaD*; n = 5; significance calculated by Mann Whitney U, p = 0.0079; two-tailed p-value. This reduction in hypervesiculation is non-significant between Δ*tamB* and Δ*mlaDΔtamB* (p = 0.0952). WT and Δ*mlaD* OMV production data previously shown in Fig 2 above.

Previous work in model diderms has inferred the function of the Mla system by exploiting other pathways that maintain lipid asymmetry, such as overexpression of the phospholipase A1 encoding gene (*pldA*) to rescue detergent hypersensitivity^7^ and radiolabelling with PagP to follow the hepta-acylation of lipid A^7, 27^. As homologues of these systems are not present in *V. parvula*, we relied on lipid extraction and comparative thin layer chromatography (TLC) to infer the function of these *mla* genes. However, to understand potential changes in lipid composition due to deletion of the Mla system, we first had to determine the nature of the major lipids present in *V. parvula* SKV38. We identified homologues of genes that may be involved in the biosynthesis of phosphatidylethanolamine (PE), phosphatidylserine (PS) and phosphatidylglycerol (PG), as well as a homologue of the recently identified hydrogenase / reductase operon required for plasmalogen biosynthesis in *Clostridium perfringens*^28^, encoded within a single gene **(Supplementary Fig S1A)**. Using TLC, matrix-assisted laser desorption/ionisation quadrupole ion trap time-of-flight (MALDI-QIT-TOF) and nuclear magnetic resonance (NMR) spectroscopy on lipid extracts from the WT, we then revealed that the major lipid species in the envelope of *V. parvula* are PE (∼68%) and PG (∼20%), along with two other uncharacterised lipids representing a further ∼9% and ∼3% of the envelope **(Supplementary Fig S1**, **Supplementary Table S1)**. NMR analyses also revealed that ∼15-35% of each lipid species present in the envelope of *V. parvula* contains a vinyl ether bond, which could result from the activity of the identified plasmalogen biosynthesis homologue **(Supplementary Fig S2)**. No cardiolipin was observed via MALDI-QIT-TOF, consistent with the absence of homologues of genes involved in the synthesis of this lipid in the *V. parvula* genome.

Both whole-cell and membrane lipid extracts from WT and Δ*mla* strains were used to perform TLC, however no differences in GPL composition were observed, as previously described in *E. coli*^7^ **(Supplementary Fig S3A)**. We posited that this lack of difference may be due to the observed hypervesiculation of Δ*mla* strains, compensating for OM lipid imbalance by shedding excess envelope material. We therefore purified OMVs produced by both the WT and Δ*mlaD* strains, performed lipid extraction and TLC, and revealed different classes of lipid species by staining with iodine vapor, Schiff reagent and phosphomolybdic acid (PMA). Across all iodine vapour-stained plates, OMVs produced by Δ*mlaD* strains display a ∼40% enrichment for PE, whilst also containing a marked reduction in another lipid species that is abundant in WT OMVs **(Fig 2C**, **Supplementary Fig S3B)**. These results suggest that Δ*mla* mutants accumulate PE in their OM, which is subsequently shed from the envelope via excessive vesicle formation. Taken together, this analysis strongly suggests that the minimal MlaEFD system of *V. parvula* is involved in retrograde GPL trafficking.

### Suppressor analysis implicates Mla and TamB in coordination of GPL homeostasis in *V. parvula*

To further strengthen the proposed function of the Mla system in *V. parvula*, we investigated the genes able to suppress detergent hypersensitivity of Δ*mla* strains. By performing random transposon mutagenesis in the Δ*mlaD* background, we identified ten chromosomal insertion mutants able to grow in SK media supplemented with 0.004% SDS and 312.5 µM EDTA. Efficiency of plating revealed that only four of these mutants could significantly rescue the Δ*mlaD* phenotype **(Fig 3A)**, of which two were of particular interest: sequencing revealed two identical Tn insertions in the downstream homologue of *tamB* **(Fig 3B)**.

**Figure 4:**
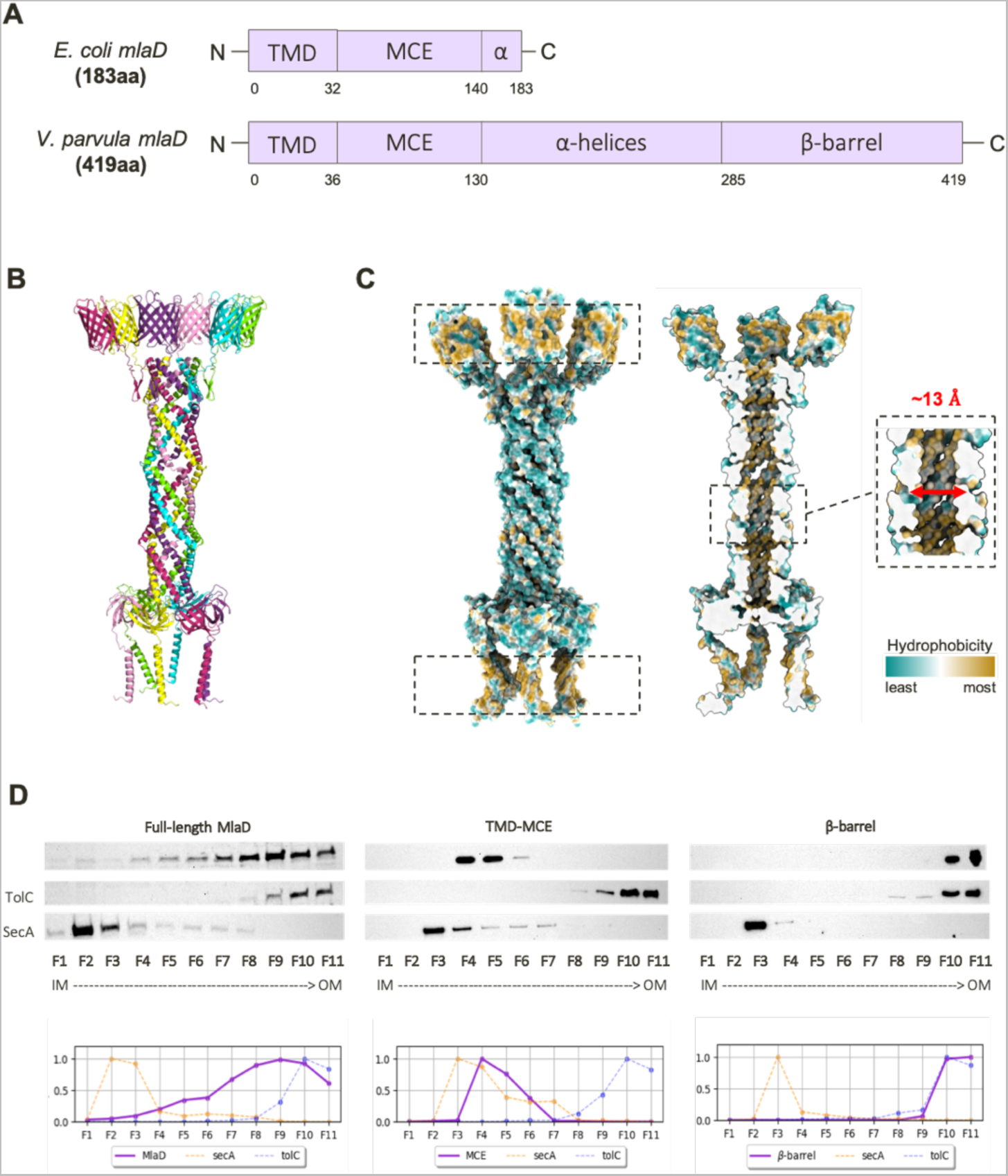
Structural modelling and experimental localisation of a transenvelope MlaD in *V. parvula*. **A) Domain comparison of MlaD in *E. coli* and *V. parvula*.** MlaD in *E. coli* consists of 183 residues and two major domains: a transmembrane domain (TMD) and an MCE domain, with a small alpha-helical region at its C-terminus. MlaD in *V. parvula* is 419 residues, consisting of a TMD, MCE domain, extended alpha-helical region and a β-barrel at its C-terminus. **B) Merged hexameric model of full-length MlaD.** Separate AF2 models of each domain of MlaD were overlapped to create a merged, full-length hexameric model of MlaD. **C) Hydrophobicity of the full-length hexameric MlaD model.** The full-length MlaD model displays two strikingly hydrophobic regions, at both the N- and C-termini, which are predicted to both be membrane-embedded domains. The tunnel of this model displays a hydrophobic interior, with a diameter of around ∼13Å. **D) Subcellular localisation of MlaD domains via sucrose-density gradient membrane fractionation.** The WT strain, and Δ*mlaD* complemented with the TMD-MCE domain / β-barrel domain with a C-ter HA-tag, were used for membrane fractionation via sucrose gradient sedimentation. Fractions were visualised with the native MlaD antisera (WT) or anti-HA antibody (MCE / β-barrel strains). The full-length protein is present in all membrane fractions, and most abundant in F7-10. The TMD-MCE construct localises to the IM, most abundant in F4-5, whilst the barrel localises almost exclusively to the final fractions, F10-11, mirroring the localisation profile of the OM control, TolC. Relative fluorescence intensity of these localisation profiles are shown below the immunoblot.

Considering the recent implications of TamB and other large AsmA-like proteins in anterograde GPL trafficking^3, 4^, we were interested to identify this gene as a suppressor of Δ*mlaD* detergent sensitivity. Indeed, we had previously identified a spontaneous suppressor of Δ*mlaD* with a SNP encoding a STOP codon in *tamB,* though the phenotype was unstable and reverted **(Fig 3B)**. We therefore generated clean, stable deletion mutants to further probe the relationship between these two genes. As with the suppressor screens, loss of *tamB* partially rescues SDS/EDTA sensitivity in Δ*mlaD* and reduces vancomycin resistance **(Fig 3C)**. Interestingly, the *tamB* mutant presents a permeability phenotype which appears to be the opposite of Δ*mla* mutants, with no change in SDS/EDTA sensitivity and an increased sensitivity to vancomycin, as previously described in *E. coli*^3^ **(Fig 3C**, **Supplementary Fig S4)**. Much like Δ*mlaD*, however, Δ*tamB* also displays a hypervesiculation phenotype, with a ∼10-fold increase in vesicle production as compared to the WT strain (Fig 3D). Strikingly, loss of *tamB* in the Δ*mlaD* background significantly reduces hypervesiculation; the Δ*tamBΔmlaD* mutant produces only ∼5-fold more vesicles than the WT **(Fig 3D)**. This rate of reduction in OMV production is comparable to the phenotype of the fully-complemented Δ*mlaEFD::mlaEFD* strain, suggesting that deletion of *tamB* is as effective at rescuing this envelope defect as reintroducing the *mla* genes. The contrasting OM permeability phenotypes of Δ*mlaD* and Δ*tamB,* together with the striking complementation of the double Δ*mlaDΔtamB* strain, hints to the antagonistic functions of these two genes in envelope biogenesis and maintenance. Overall, these results suggest that MlaEFD and TamB may represent the two major systems maintaining GPL homeostasis in the OM of *V. parvula* - likely retrograde and anterograde, respectively.

### Structural modelling suggests a transenvelope Mla system in *V. parvula*

Though phenotypic and suppressor analyses elucidated the functional role of MlaEFD, we sought to understand how three IM proteins could be structurally arranged within the envelope of *V. parvula* to facilitate GPL trafficking in the absence of MlaABC. Whilst the *mlaE* and *mlaF* homologues closely resemble those of model diderms such as the Proteobacteria, *mlaD* in *V. parvula* is over double the size of the one of *E. coli* (419 amino acids vs 183 amino acids respectively) **(Fig 4A)**. Both versions of this gene contain a transmembrane domain (TMD) that anchors the protein in the IM, and a conserved MCE domain that binds lipids^6^. However, MlaD in *V. parvula* extends with a long predicted α-helical region spanning ∼130 residues, and a predicted C-terminal β-barrel **(Fig 4A)**.

AlphaFold2^29^ (AF2) was used to predict the structure of this unusually long MlaD protein, but was unable to confidently model the elongated α-helical region, and indeed the positioning of this domain relative to the N- and C-termini of the protein **(Extended Fig 3)**. However, high confidence predictions were achieved for the TMD and MCE domain, and for the C-terminal β-barrel. This C-terminal domain displays characteristic features of a membrane-embedded β-barrel, composed of ten alternating antiparallel β-strands with a hydrophobic and neutral surface for the β-sheets, and with ‘girdles’ of negatively charged and aromatic residues which flank the top and bottom of the structure **(Extended Fig 4)**.

Considering all characterised MCE proteins to date form hexamers, we merged overlapping predicted structures of all domains of MlaD_Vp_ with hexameric stoichiometry, generating a model for the full-length protein **(Fig 4B)**. Even when generated without templates, a 6:2:2 stoichiometry of the MlaDEF complex is confidently predicted for *V. parvula,* with high similarity to the resolved structure of the MlaDEF complex from *E. coli* **(Extended Fig 3)**. Interestingly, two major models are generated for the hexameric α-helical region of MlaD_Vp_: one with a closed tunnel structure, and one with an open pore or groove with little interaction between the first and last chains of the hexamer **(Extended Fig 5**, **Extended Table 1**). We suspect that the “open groove” configuration may represent an artifact of the prediction, owing to the way AlphaFold-Multimer concatenates sequences of all chains prior to structure inference. Instead, the closed tunnel model **(Extended Fig 5)** more closely reflects the resolved crystal structure of PqiB, another MCE-domain protein from *E. coli,* which forms a closed α-helical tunnel on top of three hexameric MCE rings^6^, and the recently resolved heterohexamer of Mce1A-1F from *Mycobacterium tuberculosis*^30^. Like these resolved structures, the predicted tunnel formed by the α-helices generates a hydrophobic interior of ∼13 Å **(Fig 4C)**, supporting the possibility of a role in hydrophobic substrate transport. Further, both the N- and C-termini of the full-length model generate highly hydrophobic bands where both regions are predicted to embed within the IM and OM, respectively. The rest of the structure, which would reside in the aqueous periplasm, displays a hydrophilic exterior **(Fig 4C)**. It is interesting to note the prediction of a small external cluster of hydrophobic residues at the lowest confidence region of the tunnel – which coincides with the region that forms a groove in the higher confidence model.

**Figure 5:**
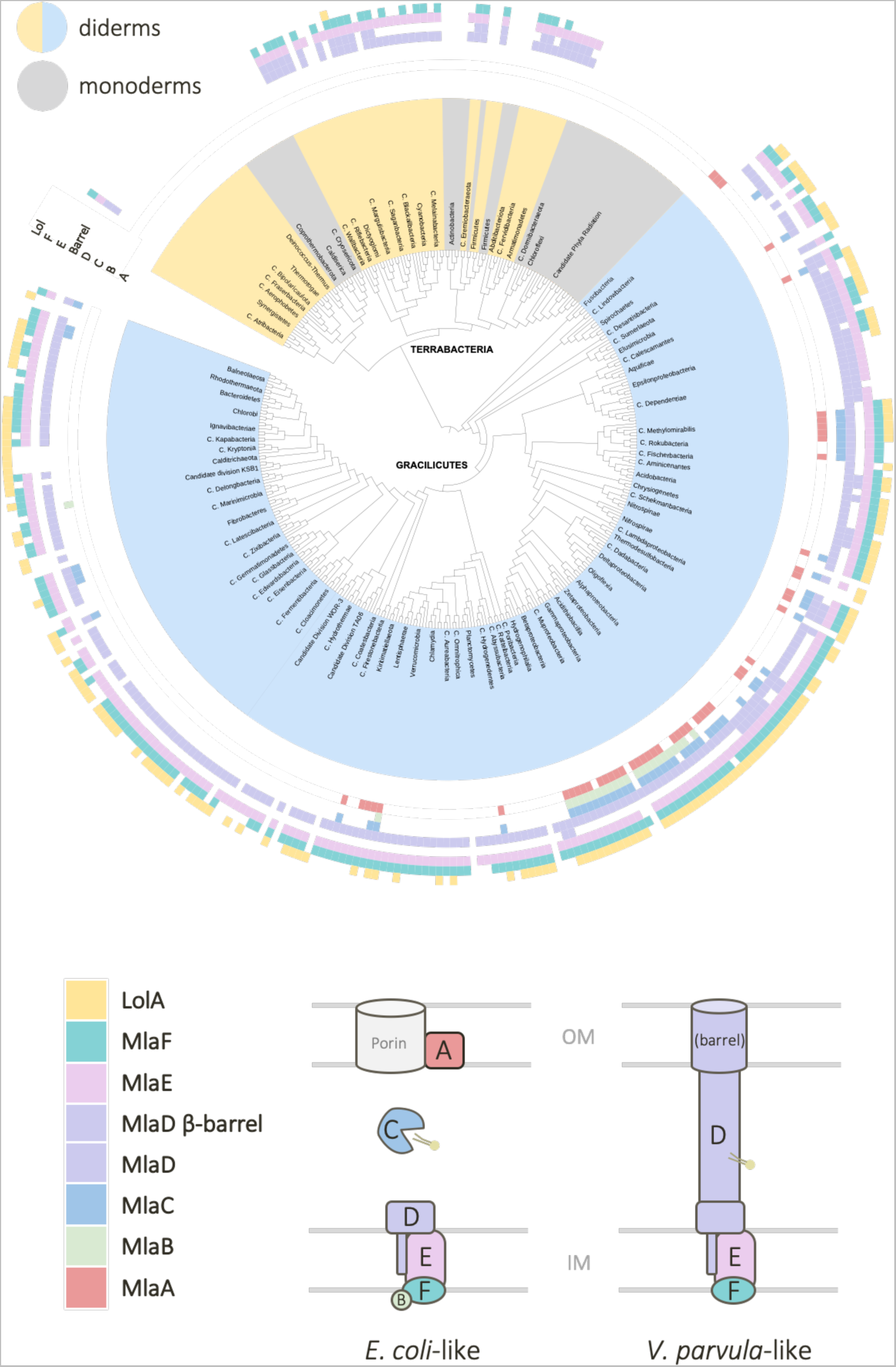
Taxonomic distribution of MlaABCDEF and LolA across Tree of Bacteria. The distribution of all six Mla components and of LolA as a proxy for the Lol system was mapped onto a reference phylogeny of Bacteria taken from ^16^ (for a more detailed version of the tree, see Extended Fig 6). In the inner ring of the tree, diderms are represented by light blue and yellow, whilst the monoderms are colored in grey. On the outer ring of the tree, each colored block corresponds to the presence or absence of the six Mla components, A-F, and of LolA. Two purple blocks are used to represent MlaD; the first block represents presence / absence of the gene itself, and the second block represents the presence / absence of the β-barrel domain. From this tree, we can see MlaEFD are widely distributed across the bacteria kingdom and were likely present in the LBCA. In contrast, MlaABC are sparsely distributed and restricted mostly to the Proteobacteria, suggesting these components evolved later (and notably the lipoprotein MlaA with the emergence of the Lol system). We also see that, in general, MlaD versions with a β-barrel tend to co-occur with the absence of MlaA and MlaC. (Yellow / Grey = Terrabacteria; Blue = Gracilicutes). Tree generated using custom made scripts and iTOL^65^. For detailed species names, see Extended Fig 6.

From these high-confidence structural predictions of a membrane-embedded C-terminal β-barrel and an IM MCE-domain hexamer, we hypothesised that MlaD from V. parvula may have domains anchored in both the IM and OM, forming a transenvelope bridge. This would remove the necessity for a periplasmic chaperone boat and OM lipoprotein, as MlaD would be capable of connecting the two membranes directly to facilitate transport. To verify this hypothesis experimentally, we localised full-length MlaD and its separate domains by performing membrane fractionation by sedimentation on sucrose density gradients. We used total membrane pellets (containing both IM and OM vesicles) obtained from WT and *ΔmlaD* strains complemented with HA-tagged versions of the TMD-MCE domain and the β-barrel. Localisation of these domains and of the full-length protein was examined via immunoblotting with our developed MlaD antisera, and the separate domains with a commercial anti-HA antibody. Whilst the full-length MlaD was distributed across almost all membrane fractions, the TMD-MCE domain construct was most abundant in fractions F4-5, similar to the localisation profile of the IM control SecA, and the β-barrel was located almost exclusively in the final few OM fractions, F9-11, mirroring the localisation of TolC **(Fig 4D)**. These results support the hypothesis that MlaD can span the periplasm, with domains anchored in both the IM and OM.

### Phylogenomic analysis reveals an ancestral core of the Mla system

The possibility of a minimal, transenvelope MlaEFD system was intriguing, and we sought to understand if this type of pathway exists in other bacterial lineages by exploring the distribution of all Mla genes. We therefore carried out an exhaustive search for homologues of the six components (MlaABCDEF) in a local databank containing 1083 genomes, representing all current phyla. To identify MlaA, MlaC, MlaD, and MlaE homologues, we used the Pfam domains PF04333, PF05494, PF02470, and PF02405, respectively. As MlaB and MlaF belong to the generic STAS-domain and ATPase families, respectively, we used MacSyFinder2^32^ to identify them when found in synteny with at least one of the other Mla components in the genome. The distribution of all six components was then mapped onto a reference phylogeny of Bacteria^16^ (**Fig 5**, **Extended Fig 6**, **Extended Table 2**).

In agreement with the fundamental role of the Mla system in maintaining OM asymmetry, Mla homologues are largely found in diderm bacteria **(Fig 5)**. Some diderm phyla in the Terrabacteria lack any identifiable homologues, possibly due to their atypical envelope architectures: the Thermotogae and their neighbor Candidate phyla Bipolaricaulota and Fraserbacteria have an OM detached at the cell poles^34^; Dictyoglomi and Deinococcus-Thermus both form rotund bodies where cell aggregates share a single OM^35, 36^; some members of Deinococcus-Thermus have lost LPS; and cryo-electron tomography of a member of Atribacteria (*Atribacter laminatus*) has recently revealed an atypical cell envelope architecture with an intracytoplasmic lipidic bilayer surrounding the nucleoid^37^. Within the three diderm lineages belonging to the Firmicutes^18^, Mla components are found in the Negativicutes and a few Limnochordia (**Extended Fig 1**), whilst they are completely absent from the Halanaerobiales (**Extended Fig 6, Extended Table 2**).

Conversely, and consistently with its presence in *M. tuberculosis,* some monoderm phyla contain MCE-domain proteins; we identified such homologues in some members of Actinobacteria, and also within the Candidate Dormibacteraeota, usually in association with a Mce4_CUP1 domain **(Extended Table 2)**. Further, our analysis suggests the MCE homologues present in these monoderm phyla were horizontally acquired from the Proteobacteria **(Extended Figs 6-8)**.

More striking is the overall distribution of MlaDEF vs MlaABC; whilst MlaDEF are widely distributed across almost all diderm phyla, with the exception of those mentioned above, the MlaABC components have a much narrower distribution, present mostly within Proteobacteria, and totally absent in diderm Terrabacteria **(Fig 5**, **Extended Fig 6**, **Extended Table 2)**.

We then focused specifically on MlaD to investigate whether other bacteria possess the longer version, similar to *V. parvula*. Surprisingly, we found that the short MlaD sequences – similar to that of *E. coli* - are restricted to taxa within Proteobacteria, whereas 85% of MlaD homologues are of the longer version (corresponding to an average of 374 residues) **(Extended Figs 6-8)**. To understand if these longer MlaD versions have a similar predicted structure to that of *V. parvula,* we screened them for the presence of β-sheet structures using BOCTOPUS2^36^ and AlphaFold2^29^. We could indeed infer the presence of a C-terminal β-barrel in 103 sequences, with a conserved fold and physicochemical properties, some containing an additional helical extension **(Extended Fig 4F, 4H)**. These long versions of MlaD with a β-barrel are widely distributed in the diderm phyla belonging to the Terrabacteria division, whilst in the Gracilicutes they are found mainly in some Proteobacteria and closely related phyla **(Extended Fig 6)**. The β-barrel is completely absent in the MlaD from the PVC (Planctomycetes, Verrucomicrobia, Chlamydia) and FCB (Fibrobacteres, Chlorobi, Bacteroidetes) groups **(Extended Fig 6**, **Extended Table 2)**. Interestingly, many of the long MlaD sequences with no identifiable complete β-barrel domain contain a disordered C-terminal region with some β- structure, often short β-hairpins, which may represent the remnants or precursors of a β*-*barrel **(Extended Fig 9)**.

Finally, to infer the origin and evolutionary history of the Mla system, we concatenated MlaD and MlaE sequences into a character supermatrix (538 leaves and 479 amino acid positions) and a maximum likelihood tree was inferred. Only markers found in a conserved cluster arrangement (MlaDEF and MlaDE) were used, due to the uncertainty of MlaF annotations. The resulting tree **(Extended Fig 7)** is consistent with the reference phylogeny of Bacteria. It shows, in fact, a clear separation between the Terrabacteria and the Gracilicutes (Ultra-Fast Bootstrap=95%), with monophyly of phyla within Terrabacteria - indicating vertical inheritance. Contrastingly, the evolutionary history of MlaDE within the Gracilicutes appears more complex, involving several duplications and horizontal transfer events **(Extended Fig 7)**.

Together, our phylogenomic analyses strongly indicate that the majority of diderms may employ a transenvelope three-component Mla system for GPL homeostasis composed of MlaDEF, with the long version of MlaD, like the one present in *V. parvula*. The six-component Mla system characterised in Proteobacteria, containing MlaABC, may therefore represent an exception, rather than the rule. Moreover, we show that this core three-component Mla system is ancestral and was present in the diderm ancestor of all Bacteria. This system was then maintained in diderm Terrabacteria and many Gracilicutes, while it underwent progressive complexification in Proteobacteria with the acquisition of additional Mla components, forming the well-characterised Mla system that we recognise in *E. coli* today.

## Discussion

In this work, we investigated a novel Mla system in the non-model diderm Firmicute *Veillonella parvula*, consisting only of MlaEFD, with no homologues of the OM lipoprotein MlaA or the periplasmic chaperone MlaC. This minimal architecture is consistent with our recent in silico studies suggesting that OM biogenesis systems present in *V. parvula* ressemble streamlined versions of those typically seen in Proteobacterial models. For example, the *V. parvula* genome encodes a homologue of BamA, but none of the associated OM lipoproteins, BamB-E; a gene encoding TamB is present, but no TamA^15, 18^. These bioinformatic findings spark a fundamental question: are these ‘missing’ components replaced by non-homologous proteins? Or do these systems function without them?

Our results show that MlaEFD likely perform a similar role in *V. parvula* as has been described in model diderms; deletion of these genes results in a loss of OM lipid asymmetry, indicated by a detergent hypersensitivity and hypervesiculation phenotype. Based on the enrichment of phosphatidylethanolamine (PE) in lipid extracts from Δ*mla* OMVs, we suggest the regular role of MlaEFD in *V. parvula* is to perform retrograde PE trafficking. We also determined that ∼15% of each lipid species present in these bacteria are in their plasmalogen form, therefore it is likely that the system shuttles plasmenylethanolamine (PlsPE) as well as typical diacyl PE.

Structural modelling and subcellular localisation helped us to understand how just three *mla* genes may perform the role of GPL trafficking in *V. parvula;* MlaEFD likely form a minimal, periplasm-spanning complex, removing the requirement for the ‘missing’ MlaABC components by directly connecting the IM and OM with a transenvelope bridge. However, future work is required to understand the stoichiometry of this protein complex, and to further explore the structure of MCE proteins across diverse bacterial lineages. Interestingly, MCE proteins in *E. coli* ^6^*, A. baumannii* ^13^ and – recently - *M. tuberculosis* ^30^ have been shown to form both hetero- and homohexamers, and for two of these structures (PqiB and Mce1A-1F), the hexameric nature of the α-helical region results in a closed tunnel that spans the periplasm, with a hydrophobic interior. These studies suggest the tunnel structures could facilitate the transport of a hydrophobic substrate and, from homology, conservation and phenotypic characterisation, the proposed substrate is a glycerophospholipid (or in the case of *M. tuberculosis,* cholesterol). However, it is difficult to understand how such a structure could support the hydrophilic head of GPLs. As phospholipid trafficking systems remain elusive even in the extensively characterised model diderms, little comparison is currently available to understand if this MlaD_Vp_ tunnel is a realistic transporter of GPLs.

So far, the only structurally characterised protein that performs retrograde GPL trafficking is MlaC, the periplasmic chaperone that can shield the acyl chains of a phospholipid whilst the head portion remains solvent-exposed ^6, 10^. This chaperone-based transport is a common theme across diderm OM biogenesis systems, from LolA shielding the tails of nascent lipoproteins ^37, 38^ to Skp / DegP and SurA protecting nascent polypeptides for delivery to the BAM system ^39^. However, our phylogenomic analysis shows that the MlaC chaperone – along with MlaA and MlaB - is only sparsely distributed in diderm bacteria. This suggests that the majority of diderms perform retrograde GPL trafficking in a very different way from *E. coli*. Considering that the long version of MlaD is actually the most common one, and that the short *E. coli-*like version of MlaD is almost entirely restricted to the Proteobacteria, it is possible that periplasm-spanning components may be a widespread method of GPL transport in many diderms. Whilst this challenges the paradigm set by extensive research in model diderms such as *E. coli* and *A. baumannii*, it is also supported by recent work in these organisms; evidence now implicates two large, periplasm-spanning AsmA-like proteins, TamB and YhdP, in anterograde lipid trafficking^3–5^. In these models, large open β-sheet structures create a hydrophobic groove that is expected to shield the acyl tails of phospholipids in a way analogous to LPS transport by the Lpt system, which employs a β- jellyroll fold, leaving the hydrophilic portion of the lipid substrate solvent-exposed^40^. Unexpectedly, two different models were generated for the tunnel region of MlaD in *V. parvula:* one with an open groove, and one closed. It is tempting to speculate that this open groove, that winds around the outside of the α-helical structure, could support the tails of the PLs in a ‘helter skelter’-like fashion, allowing the heads of the phospholipids to remain free in the aqueous environment. However, it is unlikely that six identical α-helices could form a single asymmetrical groove in its structure and, further, the hydrophobic grooves described above are composed of β-sheet structures. From the literature so far, the closed tunnel model is better supported.

Structural characterisation of *V. parvula* MlaD should help to decipher both the stoichiometry and arrangement of the MlaD complex, and its mechanism of GPL transport. This holds for the β-barrel domain which could be reminiscent of the OmpC-trimers required for the functioning of the Mla system in *E. coli* ^11, 41, 42^. Indeed, the presence of a β-barrel for substrate translocation across the OM is a common theme across other OM biogenesis systems, such as the requirement for LptD to translocate LPS to the outer leaflet of the OM^43^. However, the predicted MlaD β-barrel is too narrow to allow the passage of GPL, and a hexameric ring of β-barrels is yet to be described and seems unlikely. We posit the possibility that several of the β-barrels could fuse to form a trimeric conformation, similar to the trimeric form of TolC that results in a single, larger β-barrel^21^. Importantly, a C-terminal β-barrel was only found present in ∼10% of MlaD sequences, in both diderm Terrabacteria and a clade of Proteobacteria. In the remaining Mla homologues, we observed a variety of structures: some proteins contain disordered loops with no β-structures, whilst others contain a few β-hairpins in amongst disordered loops. While we cannot speculate at this stage on the order of the evolutionary events that led to this range of disordered, barrel-less MlaD proteins, it is interesting to consider that these unstructured regions could represent either the remnants or even precursors of a β-barrel. Considering the majority of long MlaD sequences are barrel-less, we could speculate that this structure is not important for the putative lipid importer function across bacteria. However, the recently resolved structure of *M. tuberculosis* Mce1A-1F – which do not contain complete C-terminal β-barrels - shows that each monomer contributes a single β-strand, forming a six-stranded β-barrel as a heterohexamer. Perhaps this structural organisation is repeated in other long MlaD sequences without complete barrel structures, suggesting a functional convergence across diverse bacterial lineages. The function of these barrels remains to be determined, and may differ across species; perhaps this domain is required for stability or anchoring in some bacteria, and for substrate passage in others.

Regardless of the role of the β-barrel domain, or whether the extended MlaD in *V. parvula* is capable of transporting GPLs via a winding fairground slide or a direct tunnel, the implications are the same: this mechanism is one of higher throughput than a chaperone system. Indeed, this is an important element to consider in the argument about the directionality of the Mla system. Whilst a wealth of studies support the function of Mla in retrograde GPL trafficking^7, 9, 12, 44^, and several studies argue for the anterograde direction^13, 14, 45, 46^, efficiency is a key factor to consider. In the case of anterograde GPL trafficking to facilitate cell growth, GPL flow would have to be enormously high throughput, considering the sheer volume of GPLs required to build a bilayer. In these cases, large periplasm-spanning AsmA-like proteins, which can transport several GPLs at once, would likely facilitate the high flow rate required. On the other hand, a chaperone-based system which can only offer a 1:1 ratio of protein:GPL could not accommodate this high flow rate, and likely stands as a fine-tuning mechanism – which fits with the proposed function of the Mla system, i.e. to remove surface-exposed GPLs from the OM to restore lipid asymmetry. This idea is mirrored by the Lpt system that has evolved to transport LPS into the outer leaflet of the OM, which requires a trans-envelope bridge - and not singular protein chaperones. We do not yet understand where the long and likely transenvelope MlaD fits into these scenarios; as MCE proteins are primarily implicated in importer functions, would long MlaD provide high-throughput retrograde GPL trafficking? Or could this process be bi-directional with the same transporter? Perhaps sites of hemifusion, along with accessory proteins, are necessary to control and facilitate the functioning of these trans-envelope structures, as has been suggested for the functioning of YhdP in an *mlaA\** background^5^. Intriguingly, as the long MlaD version is widely distributed across Bacteria, whatever role this protein performs is likely common across a wide diversity of lineages, waiting to be uncovered.

Furthering our understanding of GPL homeostasis in *V. parvula* is the discovery that TamB is a major suppressor of the Δ*mla* phenotype. Given the recent implications of AsmA-like proteins in anterograde GPL trafficking^3–5^, it is very interesting to discover that this protein likely performs a similar role in *V. parvula*. These findings, in combination with the recent literature, strongly advocate for contrasting but similar roles of TamB and MlaEFD in maintaining the OM in *V. parvula,* performing anterograde and retrograde GPL trafficking respectively. However, as the double Δ*mlaDΔtamB* mutant is viable, we propose that there are as yet undiscovered systems that perform lipid trafficking in *V. parvula*.

To conclude, our results reinforce the notion that the paradigms set by extensive studies within the Proteobacteria can be challenged by using non-model and phylogenetically distant bacteria. Research within diderm models such as *E. coli, A. baumannii* and *N. meningitidis* is crucial for our understanding of the functioning and virulence of many pathogens, enabling us to expand research into effective treatment options and aid identification of novel targets. However, learnings from fundamental microbiology – including those from commensals such as *V. parvula –* are often transferable, and may help us fill in the gaps regarding the structure, function and diversity of OM biogenesis and maintenance systems across the Bacterial tree of life.

## MATERIAL AND METHODS

### Bacterial strains and growth conditions

*Veillonella parvula* SKV38 was grown in SK medium (10 g/L tryptone [Difco], 10 g/L yeast extract [Difco], 0.4 g/L disodium phosphate, 2 g/L sodium chloride, and 10 ml/L 60% [wt/vol] sodium DL-lactate; described in ^47^. Cultures were incubated at 37°C in anaerobic conditions, either in anaerobic bags (GENbag anaero; bioMérieux no. 45534) or in a C400M Ruskinn anaerobic-microaerophilic station. *Escherichia coli* was grown in lysogeny broth (LB) (Corning) medium under aerobic conditions at 37°C. Where required, *V. parvula* cultures were supplemented with 20 mg/L chloramphenicol (Cm), 200 mg/L erythromycin (Ery), or 2.5 mg/L tetracycline (Tc), and *E. coli* cultures were supplemented with 100 mg/L ampicillin (Amp). 100 μg/L anhydrotetracycline (aTc) was added to induce the P*_tet_* promoter and 1mM isopropyl-β-d-thiogalactopyranoside (IPTG) was added to induce the T7 promoter. All chemicals were purchased from Sigma-Aldrich unless stated otherwise. All chemicals were purchased from Sigma-Aldrich unless stated otherwise. Primers and strains used in this chapter are listed in **Supplementary Tables S2-3**.

### Generation & Screening of Transposon mutagenesis library

A random transposon mutagenesis library was generated in *V. parvula* SKV38 WT, or the Δ*mlaD* mutant, using pRPF215, a plasmid previously used in *Clostridium difficile* that contains an aTc-inducible transposase and mariner-based transposon (Addgene 106377) ^48^. pRPF215 was transformed into *V. parvula* SKV38 by natural transformation and selected on SK agar supplemented with Cm (20μg/mL). Several independent overnight cultures of *V. parvula* SKV38-pRPF215 were diluted to OD_600_ = 0.1 in SK medium supplemented with aTc (100ng/μL) and incubated for 5 h to induce the transposase. Following induction, cultures were diluted and plated onto SK supplemented with erythromycin (200ug/μL) and aTc (100ng/μL) for selection, then incubated in anaerobic conditions for 48h. Resultant colonies were used to inoculate Greiner Bio-one polystyrene flat-bottom 96-well plates (655101) in SK supplemented with either Ery and aTc or Cm to confirm both the presence of the transposon and the loss of pRPF215. After 24h incubation, aliquots were taken from each well to inoculate fresh 96-well plates filled with SK and SK supplemented with detergents and antibiotics for the screening process. Original transposon mutagenesis plates were stored in 15% glycerol at -80°C, whilst the screening plates were incubated for 24h then assessed for growth and OM permeability defects. Mutants of interest were tested for stability of phenotype, then harvested for genomic DNA using the Wizard® Genomic DNA Purification Kit (Promega). Genomic DNA was sent for whole-genome sequencing at the Mutualized Platform for Microbiology at Institut Pasteur.

### Efficiency of plating / serial dilution assays

Sensibility to different detergents was performed on SK agar plates supplemented with detergents and antibiotics at different concentrations. Serial dilutions of strains (10^0^ - 10^-5^) were inoculated with a starting OD_600nm_ = 0.5. Plates were incubated in anaerobic conditions for 48h and *cfu* were calculated.

### Chromosomal Mutagenesis by Allelic Exchange

To generate deletion strains of *V. parvula* SKV38, site-directed mutagenesis was performed as previously described ^47^. Briefly, 1kb regions upstream and downstream of the target sequence were PCR amplified using Phusion Flash high-fidelity PCR master mix (Thermo Scientific, F548). For the selection process, three main resistance cassettes were PCR amplified with overlapping primers for the upstream and downstream regions of the target gene: the *V. atypica* tetracycline resistance cassette (*tetM* in pBSJL2), the *C. difficile* chloramphenicol resistance cassette (*catP* in pRPF185; Addgene 106367 ^49^ and the *Saccharopolyspora erythraea* erythromycin resistance cassette *(ermE).* PCR products were ligated to generate deletion cassettes via Gibson cloning or used as templates in a second PCR reaction to generate linear dsDNA with the resistance cassette flanked by the upstream and downstream sequences. These deletion constructs were transformed into *V. parvula* by natural transformation (detailed below), and its genomic integration was selected by plating on tetracycline, chloramphenicol or erythromycin-supplemented medium. Positive candidates were further confirmed by PCR and sequencing.

### Natural Transformation

Recipient strains were grown on SK agar for 48h in anaerobic conditions at 37°C. Biomass was resuspended in 1 ml SK medium adjusted to OD_600_ = 0.4 - 0.8. 10 μL aliquots were spotted onto SK agar, then 0.5 - 1 μg plasmid or 75 - 200 ng/μL linear double-stranded DNA (dsDNA) PCR product was added, using distilled water as a negative control. Following 48h incubation, the biomass was resuspended fresh SK medium, plated onto SK agar supplemented with the relevant antibiotic, and incubated for a further 48h. Colonies were streaked onto fresh selective plates, and correct integration of the construct was confirmed by PCR and sequencing.

### Growth kinetics

Overnight cultures were diluted to 0.05 OD_600_ in 200 μL SK that had previously been incubated in anaerobic conditions overnight to remove dissolved oxygen in Greiner flat-bottom 96-well plates. To maintain anaerobic conditions, a plastic adhesive film (adhesive sealing sheet, Thermo Scientific, AB0558) was used to seal the plate whilst inside the anaerobic station. The sealed plates were then incubated in a TECAN Infinite M200 Pro spectrophotometer for 24h at 37°C. OD_600_ was measured every 30m after 900 seconds orbital shaking of 2 mm amplitude.

### LPS silver staining (AgNO_3_)

#### Sample Preparation

Overnight cultures were concentrated to OD_600_ = 10 in Tricine Sample Buffer (BioRad 1610739: 200 mM Tris-HCl, pH 6.8, 40% glycerol, 2% SDS, 0.04% Coomassie Blue) and incubated at 100°C for 10min. After cooling to room temperature, Proteinase K was added to a final concentration of 1mg/mL and incubated at 37°C for 1h. 10 μL of samples were loaded onto tris-tricine gels (BioRad 4563063: 16.5% Mini-PROTEAN® Tris-Tricine Gel) and migrated in TTS buffer (BioRad 161-0744: 100 mM Tris-HCl pH 8.3, 100 mM Tricine, 0.1% SDS) for 2h at 50mA.

#### Staining

To visualise LPS profiles after gel electrophoresis separation, all development solutions were prepared immediately before use. Gels were fixed in ethanol 30%, acetic acid 10% for 1h, then washed 3 times in dH_2_O. Gels were incubated for 10m with metaperiodate 0.7%, again washed 3 times in dH_2_O, and incubated in thiosulfate 0.02% for 1m. Following a further 3 washes in dH_2_O, gels were incubated in AgNO_3_ 25mM for 10-15m, washed in dH_2_O for 15s, then incubated in development solution (K_2_CO_3_ 35mg/ml, formaldehyde 0.03% (w/v), thiosulfate 0.00125%) until bands appeared. To stop the reaction, gels were finally incubated in 4% Tris Base, 2% acetic acid for 30min, then imaged.

### Spontaneous suppressor selection

Spontaneous suppressor mutations were selected by passaging overnight cultures of Δ*mlaEFD* and Δ*mlaD*, either in liquid media or on agar, supplemented with SDS 0.004% and EDTA 312.5 μM, until growth occurred. Colonies were restreaked onto the same SDS / EDTA concentration, or directly grown in a liquid culture overnight, then genomic DNA was harvested and sent for whole genome sequencing to identify the corresponding SNPs that enabled growth of the mutants in SDS / EDTA.

### Lipid extraction and Thin Layer Chromatography (TLC)

#### Sample Preparation (Lipid Extraction)

To extract lipids, typical Bligh-Dyer extraction methods were used: briefly, overnight cultures were pelleted and concentrated to OD600 = 15 in 1mL of distilled water, then incubated on ice for 5-10min. Using glass pipettes, a monophasic mixture of 4mL methanol and 2mL chloroform were added to the resuspended cells and vortexed vigorously for 5min. Following a further 10min incubation on ice, 4ml chloroform and 2ml of distilled water were added and samples were vortexed before centrifugation at 1500 x g for 5min. This centrifugation step allowed separation of the two phases: the upper water phase, containing solutes, ions, sugars, DNA and RNA, and the lower chloroform phase, containing the lipids. The upper phase was discarded then 1mL sodium chloride solution (0.5M) was added and the samples were vortexed. Following a second centrifugation at 1500 x g for 5min, the lower phase was carefully extracted using glass micropipettes and dried in a vacuum centrifuge or under a flow of nitrogen gas (N_2_). If the initial purity was not satisfactory, a secondary sodium chloride step (and centrifugation) could be repeated. Dried lipid extracts were resuspended in 30μL chloroform.

#### Thin Layer Chromatography (TLC) and Staining

To visualise lipids, TLC plates (Silica gel 60, non-fluorescent, 0.25mm thick, Merck Millipore) were cut to a width that allowed ∼1cm between each lipid sample. Plates were pre-run in acetone before use and dried at 100°C for 30min. 10µL of each sample was loaded with 5µL glass microcapillary tubes (Sigma Aldrich) 1cm from the bottom of the TLC plate. The plate was then placed in a solvent system optimised for the separation of phospholipids by head group polarity: 65:25:4 (v/v/v) or 80:15:2.5 chloroform/methanol/water. For separation of lipids from OMVs, we used the ratio of 80:20:2.5. When the solvent front reached ∼2cm from the top of the plate, the plate was removed from the solvent system and dried at 100°C for 30m or air-dried for 10min. To reveal phospholipid species, the dried plate was incubated in a glass tank saturated with iodine vapour from iodine crystals until lipid spots were clearly visible (∼5min). After imaging this stained plate, the iodine was allowed to decolourise and the same plate was re-stained with phosphomolybdic acid (PMA, 10%), followed by vigorous heating to develop spots, or Schiff reagent (Sigma-Aldrich). Amines were detected by spraying TLC plates with 0.2g Ninhydrin (Sigma-Aldrich) dissolved in 100ml ethanol, followed by vigorous heating to develop spots. For the relative quantification of each lipid class, ImageJ was used.

### Lipid identification

#### Nuclear Magnetic Resonance (NMR)

After labile proton to deuteron exchange with CD_3_OD, phospholipids were dried under vacuum and dissolved in a mixture of CDCl_3_/CD_3_OD (2:1, v/v). The samples were then introduced into a 3mm NMR tube. Solution NMR recordings were conducted at 10 or 20 °C on Bruker AVANCE NEO 400 and 900 spectrometers equipped with a 5mm TBI and 5mm cryo-TCI probe, respectively. Standard experiments were run: 1D-^1^H, 2D-^1^H-COSY, 2D-^1^H-TOCSY, 2D-^1^H-ROESY, 2D-^1^H-^13^C-HSQC-DEPT, 2D-^1^H-^13^C-HSQC-TOCSY. After acquisition and phase correction, chemical shifts calibration on CHD_2_OD signals were performed for ^1^H (3.31 ppm) and ^13^C (49.29 ppm). The data were analysed using the TopSpin software (Bruker).

#### Mass spectrometry (MS)

MALDI-QIT-TOF Shimadzu AXIMA Resonance mass spectrometer (Shimadzu Europe, Manchester, UK) in the positive mode was used to identify the different lipids. All lipids were dissolved in CHCl_3_/CH_3_OH (1:1, v/v) and mixed with the same volume (10µL) of 2,4,6-trihydroxyacetophenone (THAP) dissolved in methanol.

### Outer Membrane Vesicle (OMV) Extraction and Purification

Overnight cultures were used to generate 1 litre cultures of the relevant *V. parvula* strains (WT SKV38 and Δ*mla* mutants). Bacterial cells were pelleted by centrifugation at 5000 x *g* for 10min, and supernatants were passed through 0.45µm and 0.22µm pore-size filters to remove any leftover cells or debris (Sarstedt). The filtered supernatant was then concentrated using Centricon Plus-70 centrifugal filters (100kDa cutoff, Millipore) centrifuged at 3,500 x *g* 4°C for 30min, 50mL at a time. ∼3mL of concentrated OMV solution was purified from 1 litre of supernatant. This OMV solution was then pelleted by ultracentrifugation at 100,000 x *g* for 3h at 4°C (Beckman Coulter Optima L-80 XP Ultracentrifuge, Type 50.2 Ti rotor). The mass of the OMV pellets was recorded and used for subsequent lipid extraction and analysis via Thin Layer Chromatography (TLC). Purified OMVs were analysed via NanoFCM confirming an average size of ∼60 nm, matching the average size of vesicles in whole cell cultures observed via both NanoFCM and cryo-electron tomography (**Supplementary Fig S5**).

### OMV quantification

Overnight cultures were adjusted for OD_600_ and diluted 1/25 and 1/50 in filtered PBS solution. The size distribution and particle concentration within the samples were analysed by nano-flow cytometry (nFCM) (NanoFCM, Inc., Xiamen, China). First, the instrument was calibrated for particle concentration using 200 nm PE and AF488 fluorophore-conjugated polystyrene beads, and for size distribution using Silica Nanosphere Cocktail (NanoFCM, Inc., S16M-Exo and S17M-MV). The flow rate and side scattering intensity were then calculated based on the calibration curve, and used to infer the size and concentration of the large and small events present in each sample using the NanoFCM software (NanoFCM Profession V2.0). All samples were diluted to ensure the particle count fell within the optimal range of 2000-12,000/min, and all particles that passed by the detector during time intervals of 60s were recorded for each test.

### Membrane fractionation by sucrose gradient sedimentation

1.5L overnight cultures were pelleted at 5,000 x g for 15min and resuspended in ∼10mL HEPES buffer. Resuspended pellets were supplemented with 100µL benzonase and a small amount of lysozyme prior to lysis by two passages through a high-pressure French press (French Press G-M, Glen Mills) homogenizer at 20,000 psi. Following lysis, cell debris was pelleted at 15,000 x g for 90min. Resulting supernatant was spun in an ultracentrifuge (Beckman Coulter Optima L-80 XP Ultracentrifuge, Type 50.2 Ti rotor) at 35,000 x g for 90min to obtain the membrane pellet, containing both IM and OM vesicles. Membrane pellets were resuspended in 750µl of HEPES buffer using a Dounce homogeniser, then 500µL was loaded onto a discontinuous sucrose gradient comprising a bottom 2mL layer of 51%, a 3ml layer of 37%, a 4mL layer of 31%, and a 2.5mL top layer of 23% (w/w) sucrose. Sucrose gradient columns were centrifuged at 35,000 x g for 40h (Beckman Coulter Optima L-80 XP Ultracentrifuge, Type SW41 Ti rotor). At the end of the sucrose gradient sedimentation, 1mL by 1mL was extracted from the top of the gradient column to the bottom, resulting in 1ml membrane fractions.

### Structural modelling

For the prediction of protein domains, SMART (Simple Modular Architecture Research Tool)^50^ and I- TASSER^51^ were used. AlphaFold2^29^ was used to generate structural predictions of the Mla proteins in *V. parvula*. To specifically analyse the presence of β-barrels within our target sequences, the BOCTOPUS2 database was used^36^. As another transmembrane site–determining web-based tool, TMHMM 2.0 (Transmembrane Hidden Markov Model) was also used to analyse the transmembrane sites of proteins^52^.

All structure predictions with AlphaFold2 were performed through the ColabFold^53^ implementation. Output models of monomeric full-length MlaD (5 models) were ranked according to the pLDDT score. MlaEFD_1-130_ predictions (10 models) were obtained without structural templates and sorted by predicted TM-scores (*pTM*) since it predates the release of AlphaFold-multimer^54^.MCE, α-helical and β-barrel domains in Chimera^55^ and later refined with Modeller^56^ to correct the topology of the rotated hinges. MlaEFD_1-130_ predictions (10 models) were obtained without structural templates and sorted by predicted TM-scores (*pTM*) since it predates the release of AlphaFold-multimer^54^. To impose the 2:2:6 stoichiometry of the complex, 2 copies of the MlaE and MlaF sequences were concatenated together with 6 copies of the MlaD_1-130_ sequence. Model of the hexameric full-length MlaD complex was obtained by stitching together predictions for the MlaD_1-130_, MlaD_36-263_ and MlaD_239-419_ domains, via structural superimposition of the overlapping regions. Predictions for the MlaD_36-263_ and MlaD_239-419_ domains were first obtained using the AlphaFold2-multimer v2.3 parameters with templates found in the PDB70 database^57^ and ranked by *multimer* score^54^.

Full-length monomeric models of long MlaD sequences were obtained with AlphaFold2 as described above. Secondary structure class of all positions in the best model (highest plDDT) for each sequence were obtained using DSSP^58^ and PROSS^59^. To automatically assess the presence of residues in β-strand conformation after the MCE domain, the start and end positions of the MCE domain in each sequence was obtained using HMMSEARCH^60^ with the “MlaD protein” pfam profile (PF02470).

### Phylogenetic Analysis

We assembled a databank of 1,083 genomes representing all bacterial phyla present at the National Center for Biotechnology (NCBI). We selected three species per order for each phylum. The number of genomes per phylum therefore reflects their taxonomical diversity (Extended Data Table 2). We chose preferably genomes from reference species and the most complete assemblies. We then queried this databank for the presence of MlaA, MlaB, MlaC, MlaD, MlaE, MlaF, and TamB, using the pfam domains PF04333, PF05494, PF02470, PF02405 and PF04357 respectively. The alignments of protein families NF033618 (MlaB) and PRK11831 (MlaF) were downloaded from NCBI (Conserved Domain Database) and used to build HMM profiles using HMMBUILD from the HMMER3.3.2 package^60^ MlaA, MlaC, MlaD, MlaE and TamB homologs were searched using HMMSEARCH from the HMMER3.3.2 package. As MlaB and MlaF belong to large protein families, we used MacSyFinder2^31^ to identify their occurrences when they are in genomic synteny with at least one of the other components (*mlaACDE*) of the Mla system, with no more than five genes separating them. This also allowed the identification of the occurrences of *mlaACDE* when they were in genomic cluster. All the retrieved hits were curated using functional annotations, domains organization, alignments, and phylogeny. We identified 1119 MCE-domain containing sequences (Extended Data Table 2). Among these, homologues of PqiB and LetB (containing more than one MCE domain) were identified only within Proteobacteria and closely related phyla.

For each protein, curated sequences were aligned with MAFFT v7.407^61^ with the L-INS-I option and trimmed using BMGE-1.12^62^ with the BLOSUM30 substitution matrix. Maximum likelihood trees were generated for each protein using IQ-TREE.2.0.6^63^ with the best evolutionary model assessed with ModelFinder^64^ according to the Bayesian Information Criterion, and ultrafast bootstrap supports computed on 1000 replicates of the original dataset. Finally, the presence or absence of each protein was mapped onto a reference tree of Bacteria taken from^16^. All tree figures were generated using custom made scripts and iTOL^65^.

## STATISTICS AND REPRODUCIBILITY

Data are presented as the mean ± standard deviation (SD) and with individual data points from at least three independent biological experiments. Statistical analyses were performed using Prism 9.5.0 (GraphPad Software Inc.). The distribution of data was analysed by Shapiro-Wilk test and their variance using Bartlett test. Since some of the data generated did not follow a normal distribution or had two high variance we performed non-parametric two-tailed Mann-Whitney test. P-values <0.05 were defined as the level of statistical significance.

## DATA AVAILABILITY STATEMENT

The authors declare that all data supporting the findings of this study are either available within the paper and its supplementary information files or, otherwise, are available from the corresponding author upon request.

## ACKNOWLEDGEMENTS

We gratefully thank Noa Guzzi, and members of the UGB and EBMC units, for their assistance in generating the transposon mutant libraries used in this study. We also thank Robert Smith for his help in scaling up large anaerobic cultures for *V. parvula.* We thank Pierre-Henri COMMERE for his help with the use of the NanoFCM machine. We also thank Philippe Delepelaire for generously providing us with anti-SecA and anti-TolC serum, and Jean-Michel Betton for his help in developing the membrane fractionation protocol. This work was supported by funding from the French National Research Agency (ANR) (Fir-OM ANR-16-CE12-0010) and (OM-LipAsy-CE44-008), by the French government’s Investissement d’Avenir Program, Laboratoire d’Excellence “Integrative Biology of Emerging Infectious Diseases” (grant n°ANR-10-LABX-62-IBEID) and by the Fondation pour la Recherche Médicale (grant DEQ20180339185). KG was supported by the Pasteur Paris University program (PPU). B-Beaud was supported by a MENESR (Ministère Français de l’Education Nationale, de l’Enseignement Supérieur et de la Recherche) fellowship. Financial support from the IR INFRANALYTICS FR2054 for conducting the research is gratefully acknowledged. The authors acknowledge the IT department at Institut Pasteur, Paris, for providing computational and storage services (TARS cluster).

## AUTHORS CONTRIBUTIONS STATEMENT

KG performed all molecular biology and microbiology experiments with the assistance of BA. B-Beaud developed and optimised membrane fractionation protocols for *Veillonella parvula* and lipid extraction together with KG. NT performed all evolutionary and phylogenetic analyses. B-Bardiaux performed the structural modeling analysis. YR, XT, ZC and YG performed lipidomic analyses. JMG provided lab facilities. NIP provided lab facilities and expertise for protein purification, assisted by ML. CB and SG supervised the study. KG, CB, and SG wrote the paper with contributions from NT, B-Bardiaux and JMG. All authors contributed to the final version of the manuscript.

## Extended Data

**Extended Figure 1:**
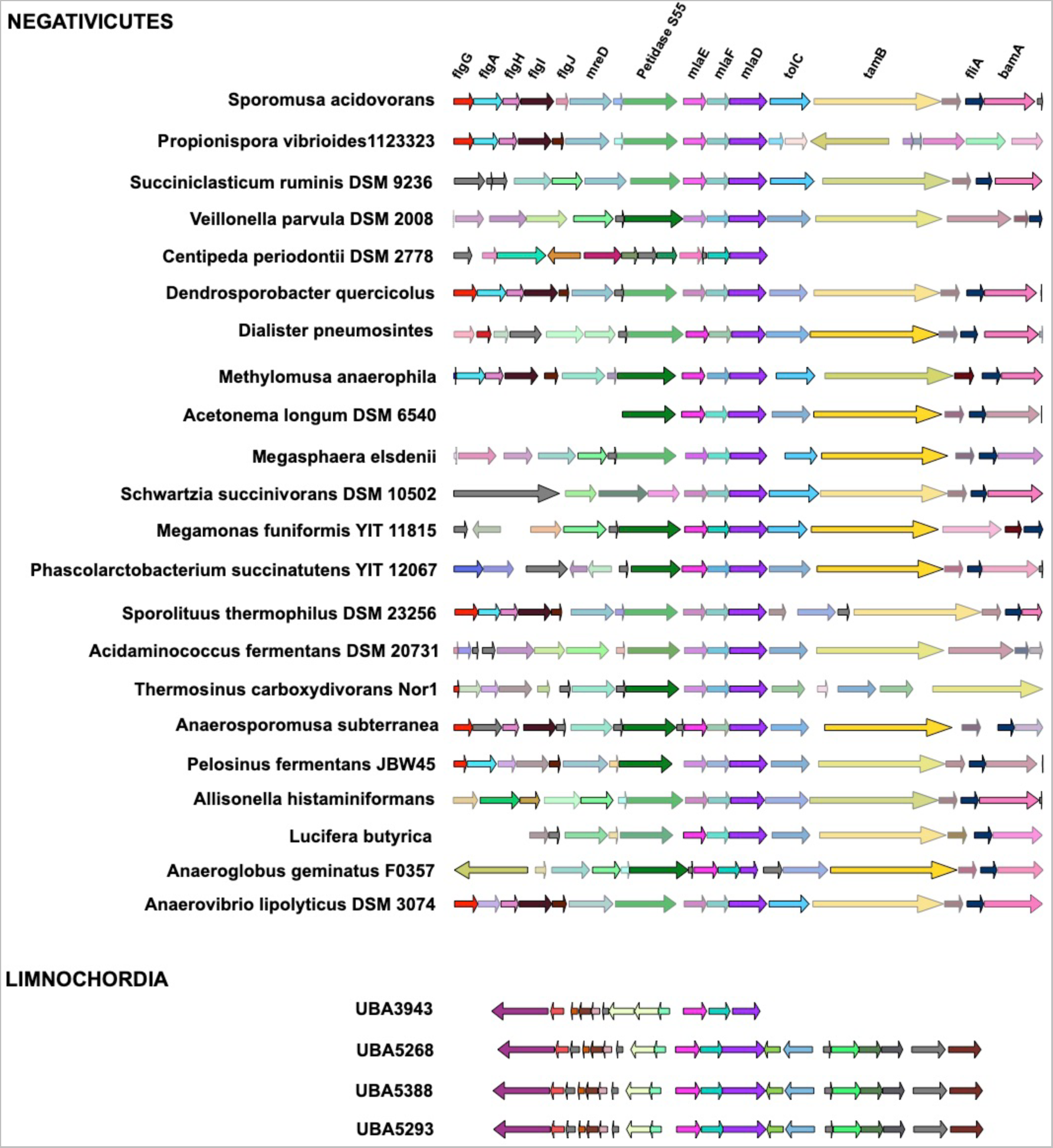
Conserved synteny of *mla* genes with *tamB* and *tolC* in the Negativicutes, and presence of *mla* genes in Limnochordia. Within the Negativicutes, the three *mla* homologues, *mlaEFD*, are directly followed by a homologue of *tolC* and a homologue of *tamB*. This synteny is not conserved within the other diderm Firmicutes; no *mla* homologues were identified in the Halanaerobiales, whilst *mla* genes in the Limnochordia are spare and not in synteny with *tolC* and *tamB* homologues.

**Extended Figure 2:**
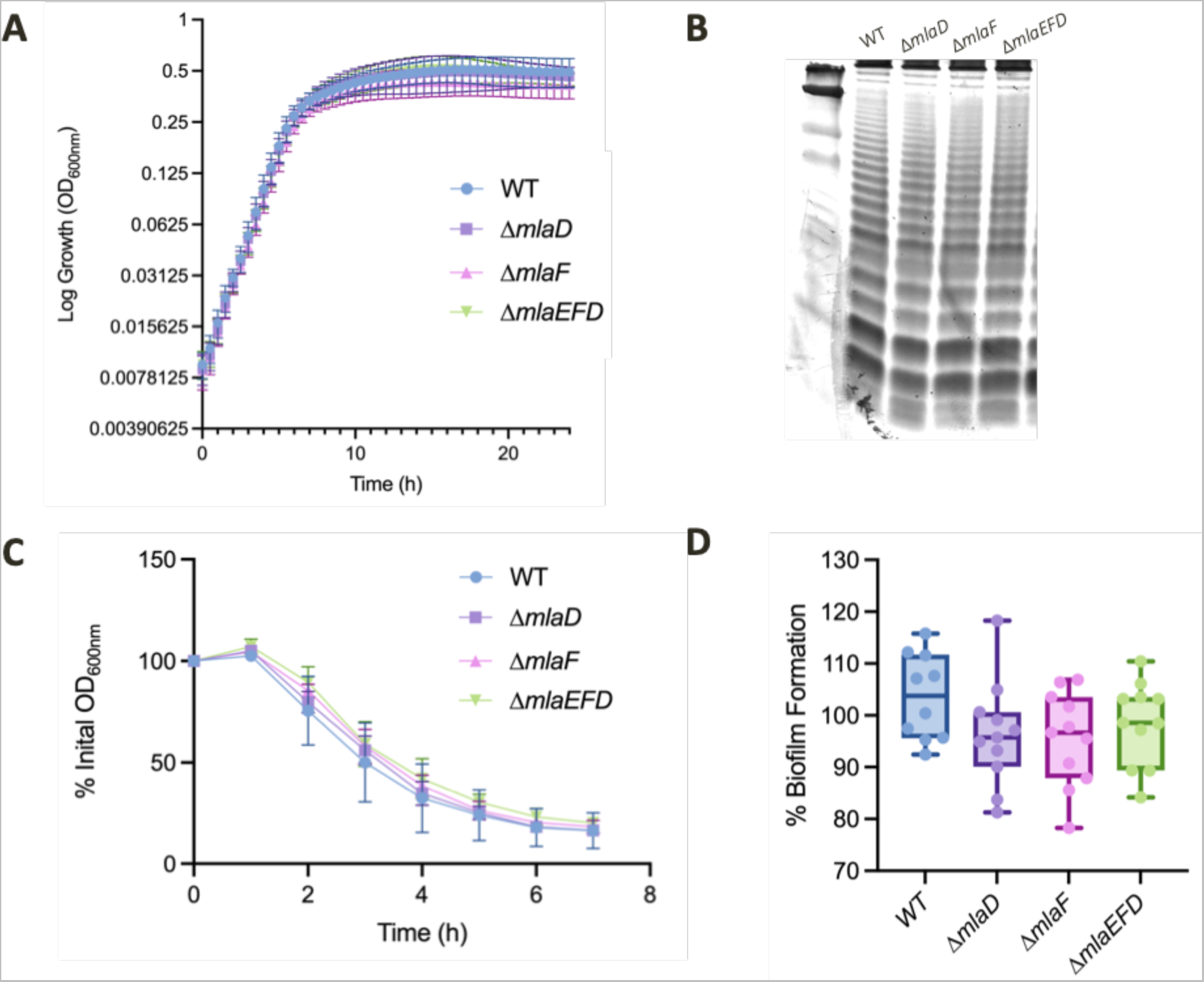
Phenotypes unchanged by deletion of *mla* genes. **A) Comparison of growth kinetics of WT and *Δmla* mutants**. All strains were assessed for growth kinetics via OD_600_ measurements via TECAN spectrophotometer (see methods). Δ*mlaD, ΔmlaF* and Δ*mlaEFD* were found to display similar growth rates to the WT, with no defects. At least 3 biological replicates and 3 technical replicates were performed per strain. **B) Comparison of LPS profiles.** LPS was extracted from WT and all Δ*mla* cultures, in triplicate, normalised by OD_600_, then observed with silver staining. All strains display similar LPS profiles, in both banding pattern and quantity. **C) Comparison of autoaggregation capabilities.** OD_600_ was recorded for WT and Δ*mla* strains once an hour for a period of seven hours. All strains display a similar decrease in OD_600_ over time, suggesting a similar capability of autoaggregation. At least 3 biological replicates and 3 technical replicates were performed per strain. **D) Comparison of biofilm formation.** Biofilm formation was assessed via crystal violet staining of biofilms formed over 24h in 96-well plates. Δ*mla* strains display a small but non-significant decrease in biofilm formation capability, as assessed by a decrease in fluorescence signal at A575nm. Due to variability in this technique, 10 biological replicates with 3 technical replicates each were tested per strain.

**Extended Figure 3:**
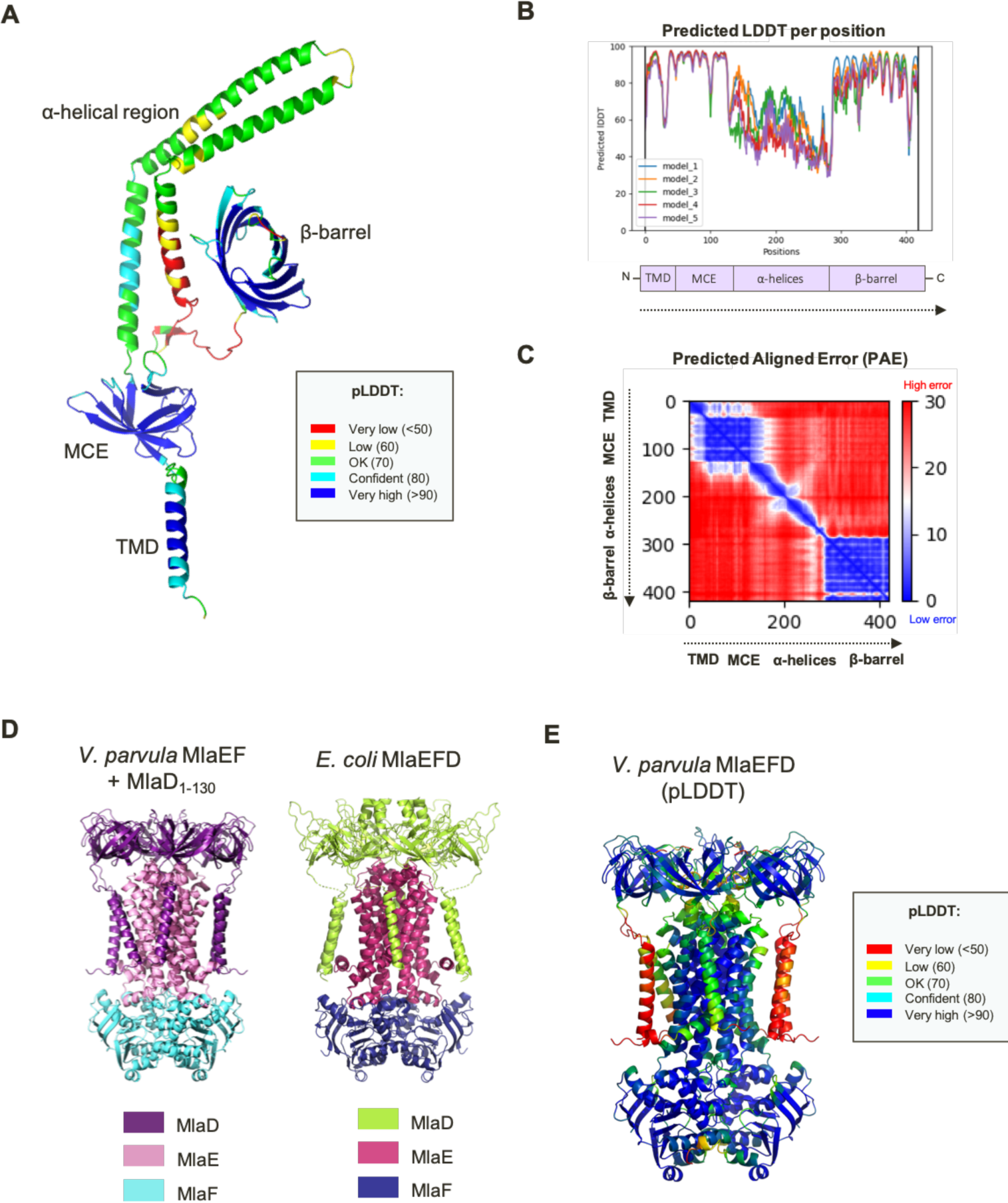
AlphaFold2 modelling of MlaD in *V. parvula*. **A) Predicted structure of full-length MlaD**. When the whole sequence is processed by AlphaFold2, the TMD, MCE and β-barrel domain are of very high confidence, and the α-helical domain is of lower confidence. The confidence rating is indicated by Predicted Local Distance Different Test (pLDDT) colouring. **B) Predicted Local Distance Difference Test (pLDDT) per position of full-length MlaD.** Graphical depiction of the pLDDT per position of the full-length models shows low confidence for the α-helical region, and high confidence for the TMD, MCE and β-barrel domains. **C) Predicted Aligned Error (PAE) of full-length MlaD.** Graphical depiction of the PAE shows the confidence in the relative positioning of the domains. The relative positioning of main chain atoms within the TMD and MCE domains and within the connecting α-helical region are low error, suggesting the predicted local conformation of these individual domains to be likely. However, the high predicted alignment error of atoms across these domain regions indicates that the global conformation of full-length MlaD is likely inaccurate. **D) Modelling of MlaEFD complex in *V. parvula* with a 2:2:6 stoichiometry.** MlaEF modelled as dimers in complex with hexameric MlaD (residues 1-130) closely resemble the resolved structure of the MlaEFD complex in *E. coli.* **E)** pLDDT per position of MlaEFD in *V. parvula* shows high confidence with the exception of 2 pairs of TM helices.

**Extended Figure 4:**
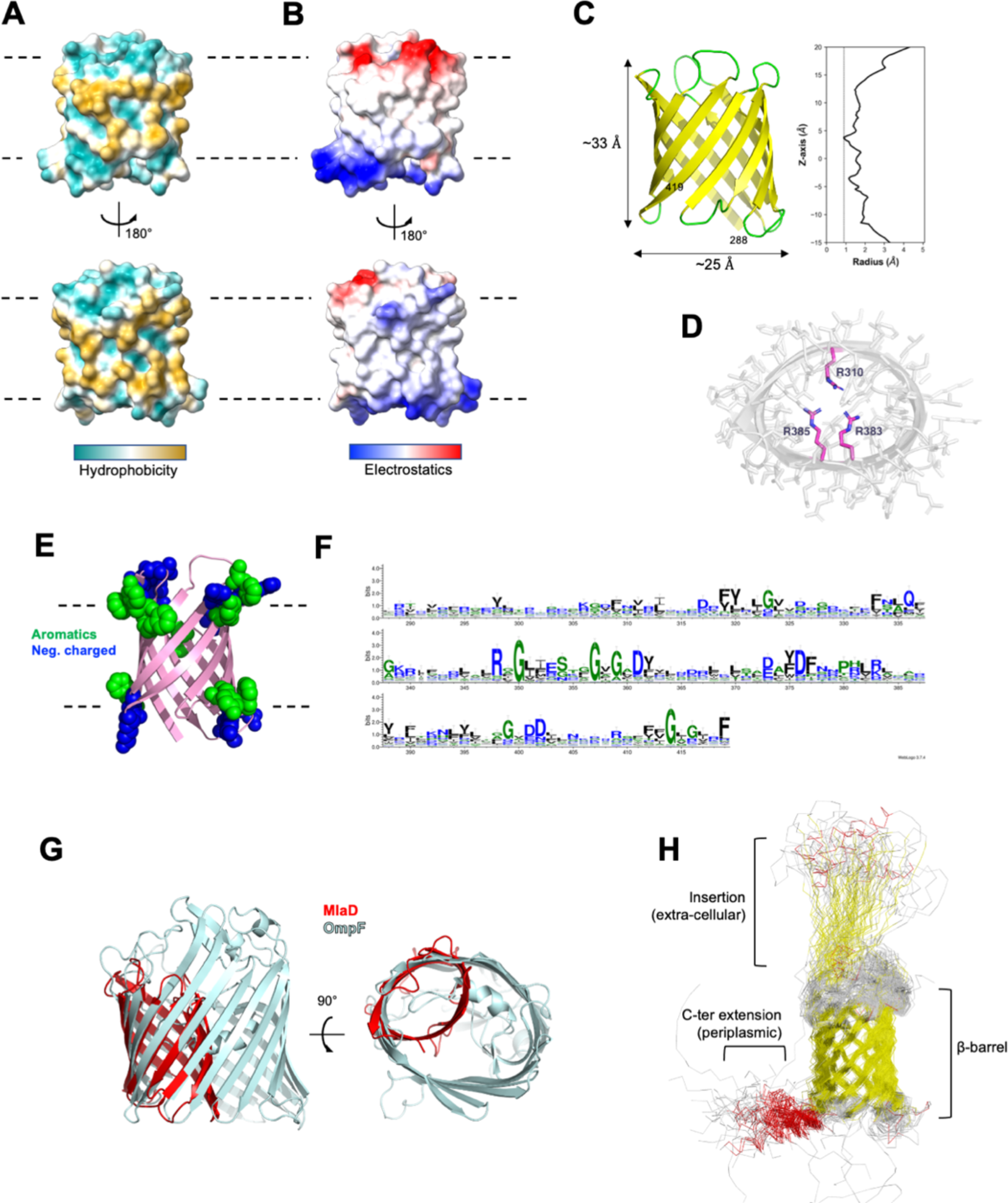
The C-terminal domain of MlaD_288-419_ folds as a membrane β-barrel. **A) Hydrophobicity potential of the MlaD C-terminal β-barrel surface**. The bottom view is rotated 180°. Dotted lines represent the possible position of GPL polar heads when inserted in a lipid bilayer. **B) Electrostatics potential of the MlaD C-terminal β-barrel surface.** The bottom view is rotated 180°. **C)** Dimensions of the β-barrel (right) and radius profile (calculated using the program HOLE) along the Z-axis (membrane normal). The height of the barrel matches the thickness of a lipid bilayer. **D)** Position of side-chains of R310, R383 et R385 at the constriction point of the β-barrel. **E)** Rings of surface accessible negatively charged and aromatics residues at each side of the β-barrel. **F)** Weblogo of MlaD C-terminal β-barrel sequences coloured by hydrophobicity (Hydrophilic in blue, neutral in green and hydrophobic in black). **G)** Superimposition of the *V. parvula* MlaD C-terminal β-barrel model (red) onto the *S. marcescens* OmpF structure (light cyan). Its small diameter, compared to large porins such as OmpF, and very narrow inner radius (∼1 Å) would likely prevent the passage of any small molecule. **H)** Superimposed ensembles of MlaD C-terminal β-barrel models for long MlaD sequences predicted with AlphaFold2. Helices are coloured in red and β-strands in yellow. Insertions in the barrel and C-terminal helical extensions found in some sequences are labelled.

**Extended Figure 5:**
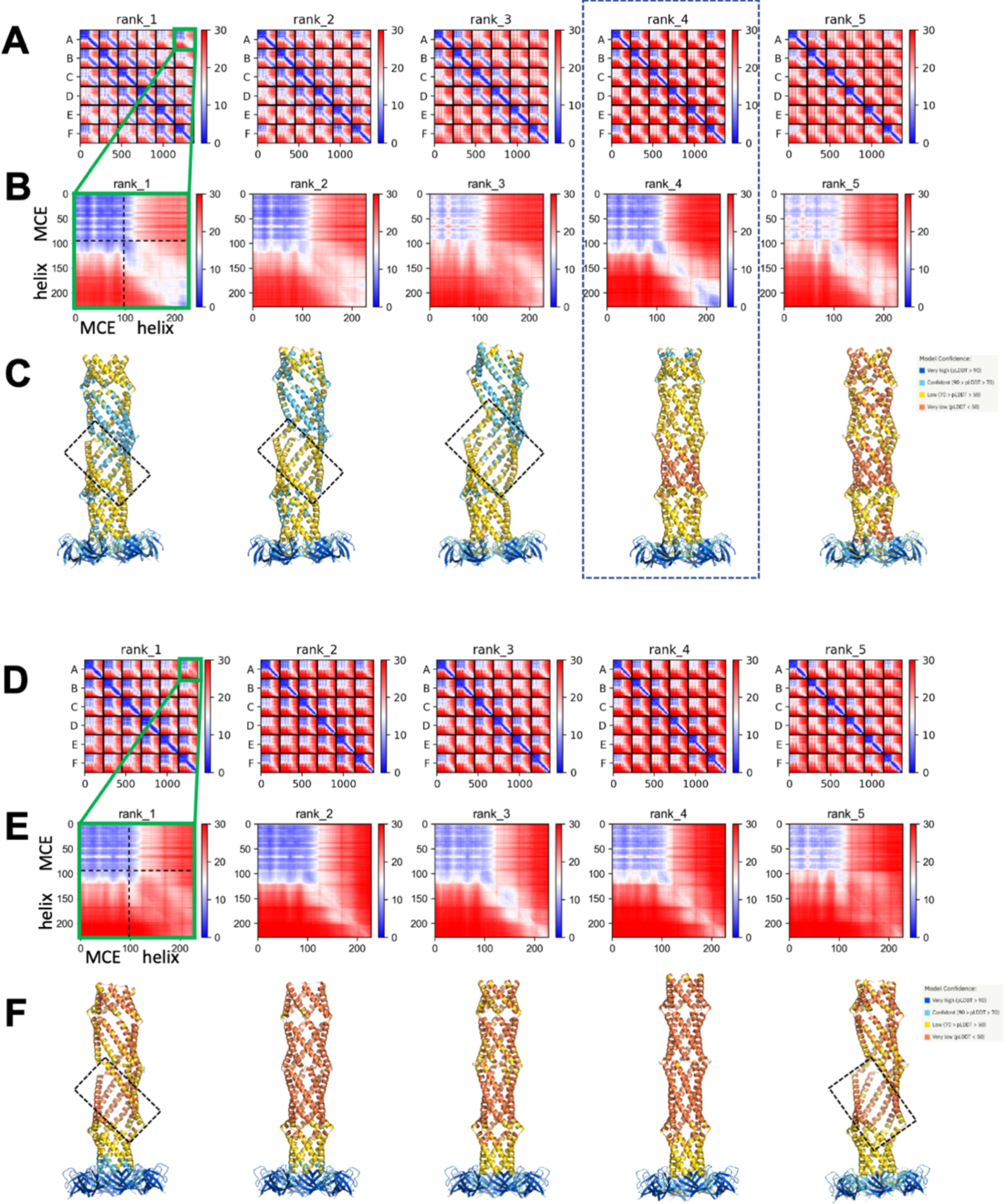
AlphaFold2 models for the alpha-helical domain of MlaD_36-263_ in hexameric configuration. **A)** Predicted Aligned error (PAE) for the 5 ranked models obtained with AlphaFold-Multimer v2.2. The regions corresponding to each chain in the hexamer (from A to G) are labeled. **B)** PAE between the first (A) and last (G) chain for the 5 ranked models obtained with AlphaFold-Multimer v2.2. The regions corresponding the MCE and helical domains are labeled. **C)** Models obtained AlphaFold- Multimer v2.2. colored by predicted local distance difference test (pLDDT). **D)** Predicted Aligned error (PAE) for the 5 ranked models obtained with AlphaFold-Multimer v2.3. The regions corresponding to each chain in the hexamer (from A to G) are labeled. **E)** PAE between the first (A) and last (G) chain for the 5 ranked models obtained with AlphaFold-Multimer v2.3. The regions corresponding the MCE and helical domains are labeled. **F)** Models obtained AlphaFold-Multimer v2.3 colored by predicted local distance difference test (pLDDT).

**Extended Figure 6:**
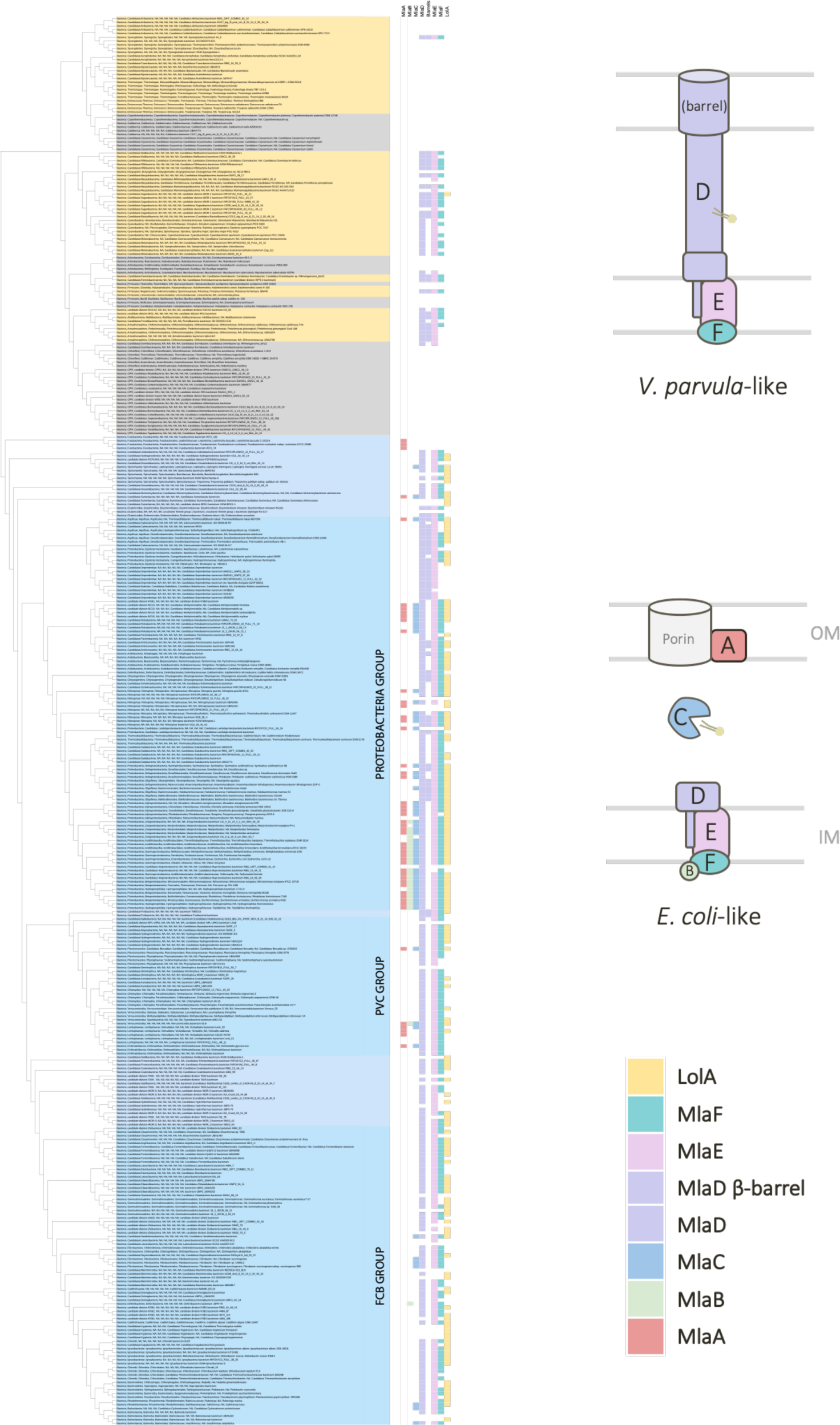
Taxonomic distribution of Mla components (and LolA) with species names. This figure is adapted from main text Fig 5 to include detailed species names. All other aspects of the figure are the same. Presence of absence of Mla component indicated in red (MlaA), green (MlaB), blue (MlaC), purple (MlaD), pink (MlaE), turquoise (MlaF) and LolA is indicated in yellow. (Yellow / Grey = Terrabacteria; Blue = Gracilicutes). Tree generated using custom made scripts and iTOL^1^.

**Extended Figure 7:**
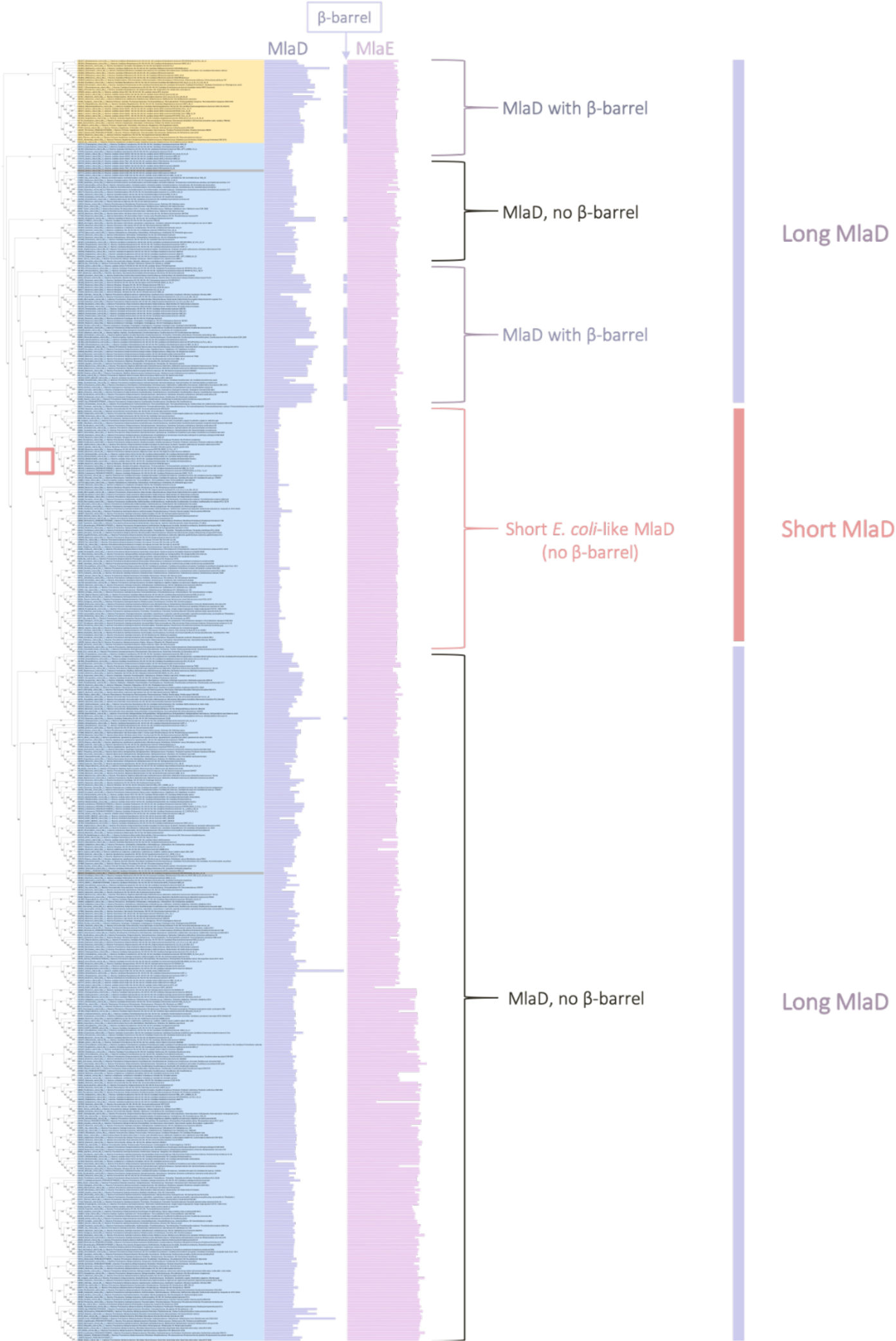
Phylogeny concatenation of MlaE with the three types of MlaD: short, long and long with β-barrel. Sequences of MlaE and MlaD were concatenated into a character supermatrix (538 sequences and 479 amino acid positions) and a maximum likelihood tree was inferred. The length of MlaD sequence is indicated by the length of the purple bars, and the presence / absence of the β-barrel is depicted with a small purple at the end of the sequence. MlaE sequences are represented in pink. We clearly see from this analysis that the majority of MlaD sequences are long, and that the short MlaD sequences are restricted mostly to the Proteobacteria. We also see a clear separation between the Terrabacteria (yellow) and the Gracilicutes (blue), with monophyly of phyla within the Terrabacteria, indicating vertical inheritance of these Mla genes. Highlighted in the red box is the branch at which loss of this elongated form of MlaD seems to have occurred.

**Extended Figure 8:**
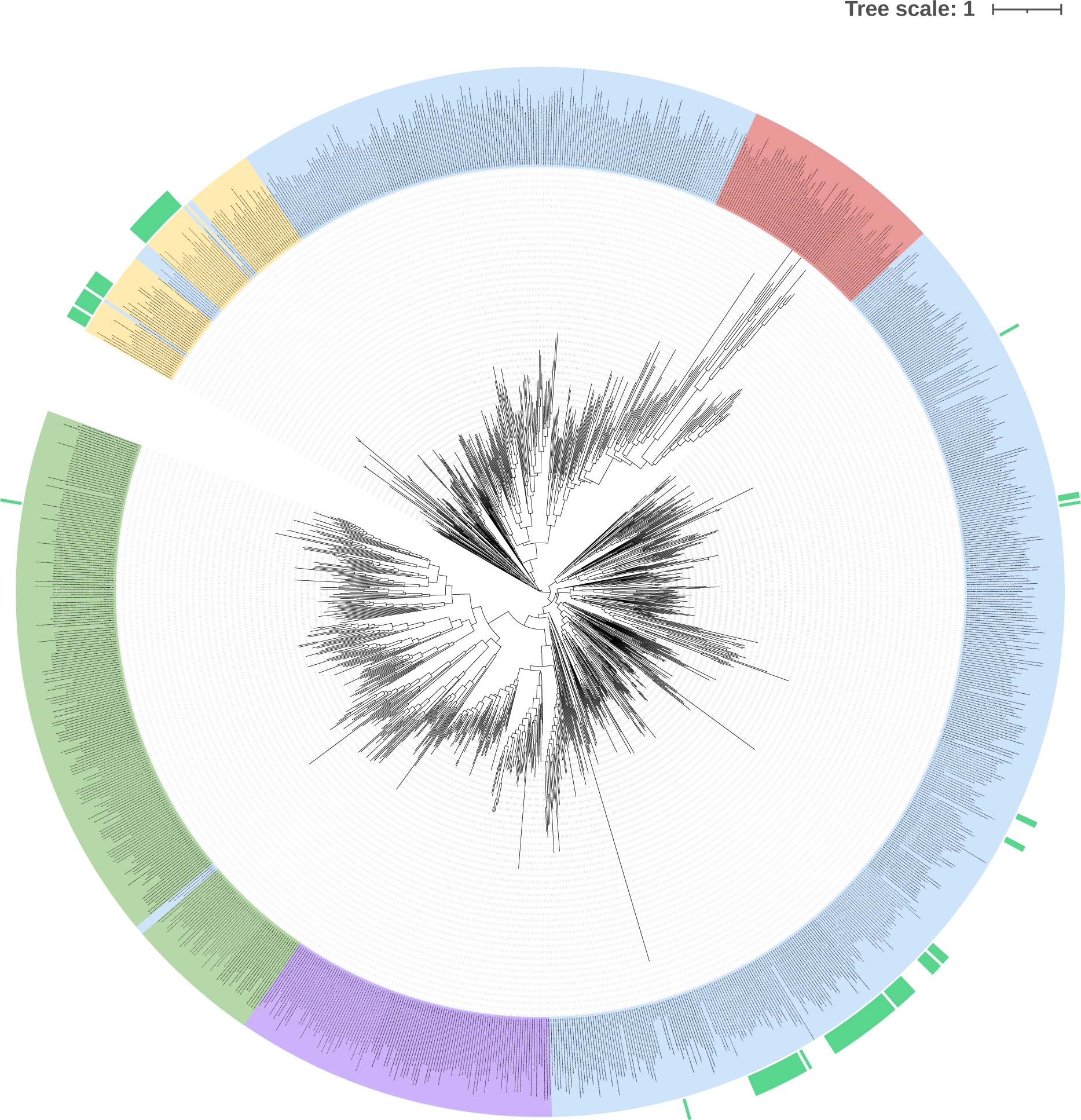
Distribution of MCE-containing sequences across Bacteria. Maximum likelihood tree built from an alignment of 1119 sequences and 194 amino acid positions. The scale bar corresponds to the average number of substitutions per site. MlaD sequences are marked in green for Actinobacteria and C. Dormibacteraeota, yellow for other Terrabacteria and blue for Gracilicutes. We highlighted the short version of MlaD in purple, and PqiB and LetB sequences in red. The presence of a β-barrel is highlighted by a green bar at the outer ring of the tree.

**Extended Figure 9:**
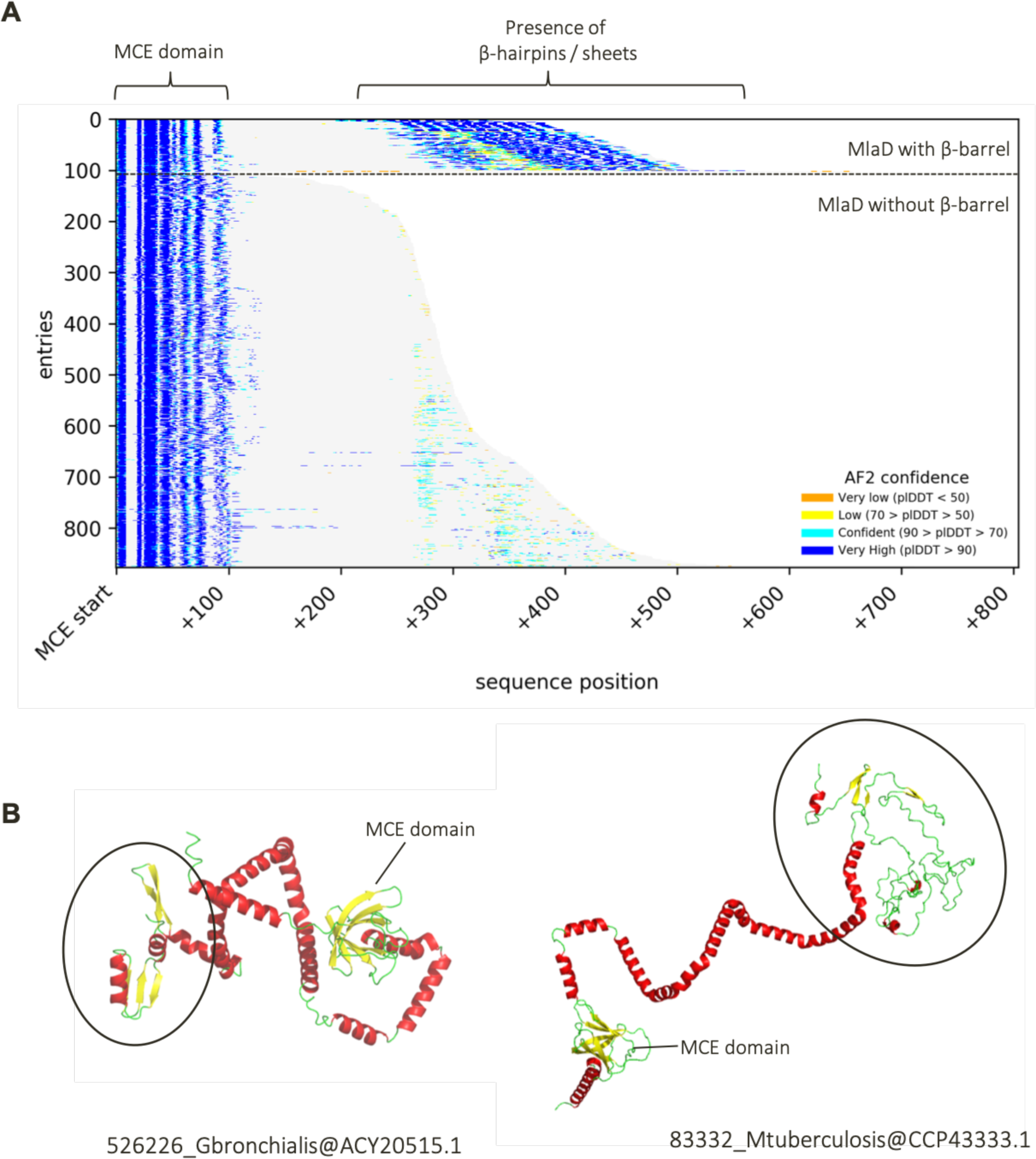
Presence of disordered β-structures in long MlaD sequences with no β-barrel. **A) Presence of β-structures in long MlaD sequences with no β-barrel**. The start of the sequence represents the presence of the MCE domain, to align all long MlaD sequences. The total length of each sequence is represented by the grey shaded area. Later, in some of these long sequences, we see the presence of β-hairpins / sheets, but no real folded β-barrel domain. For comparison, we also show the ∼100 sequences of long MlaD that do possess a predicted folded β-barrel. Predicted Local Distance Difference Test (pLDDT) values, corresponding to the confidence of these predictions, are coloured and labelled in the key above. **B) AlphaFold modelling of long MlaD sequences with no β-barrel.** As seen from these two predicted structures, these long MlaD sequences do not possess a β- barrel, but do possess unstructured, disordered regions at their C-termini that contain fragments of β- hairpins / sheets.

**Extended Table 1:** Statistics of AF2-Multimer models generated for MlaD36-263 hexamers

|  | AF-multimer v2.2 |  |  |  |  | AF -multimer v2.3 |  |  |  |  |
| --- | --- | --- | --- | --- | --- | --- | --- | --- | --- | --- |
| Model rank | 1 | 2 | 3 | 4 | 5 | 1 | 2 | 3 | 4 | 5 |
| Open/Closed | Open | Open | Open | **Closed** | **Closed** | Open | **Closed** | **Closed** | **Closed** | Open |
| pLDDT | 75.38 | 75.57 | 73.79 | 68.07 | 62.59 | 68.23 | 64.88 | 65.96 | 64.24 | 64.73 |
| pTM | 0.56 | 0.55 | 0.50 | 0.47 | 0.46 | 0.58 | 0.54 | 0.53 | 0.49 | 0.45 |
| ipTM | 0.53 | 0.52 | 0.48 | 0.44 | 0.43 | 0.56 | 0.52 | 0.50 | 0.46 | 0.42 |
| multimer | 0.54 | 0.53 | 0.48 | 0.45 | 0.44 | 0.56 | 0.52 | 0.51 | 0.47 | 0.43 |
| Contacts* | 187/**20** | 180/**0** | 190/**0** | 139/**86** | 126/**83** | 185/**3** | 101/**99** | 143/**151** | 134/**126** | 154/**5** |
| PAE < 8 Å<sup>#</sup> | 3431/**72** | 1378/**0** | 1633/**0** | 350/**102** | 16/**0** | 180/**0** | 95/**24** | 43/**0** | 4/**0** | 0/**0** |

\* Number of contacts ( $C\beta-C\beta < 8 \text{ \AA}$ ) for the helical regions (MlaD<sub>132-263</sub>) between neighboring chains. The first number is the average of all $i, i+1$ pairs of chains except between the 1<sup>st</sup> and the last chain ( $i$ being the index of the chain in the hexamer). The second number (in bold) is between the 1<sup>st</sup> and the last chain in the hexamer.
<sup>#</sup> Number of residue pairs with a Predicted Aligned Error (PAE) < 8 Å for the helical regions (MlaD<sub>132-263</sub>) between neighboring chains. The first number is the average of all $i, i+1$ pairs of chains except between the 1<sup>st</sup> and the last chain ( $i$ being the index of the chain in the hexamer). The second number (in bold) is between the 1<sup>st</sup> and the last chain in the hexamer.

**Extended Table 2:** Distribution and accession numbers of Mla components homologs among 1083 genomes representing the bacterial diversity.

The presence of MlaB and MlaF is only indicated when they are in cluster with at least one of the other components of the Mla system.

Table attached separately.

## Supplementary Data

**Supplementary Figure S1:**
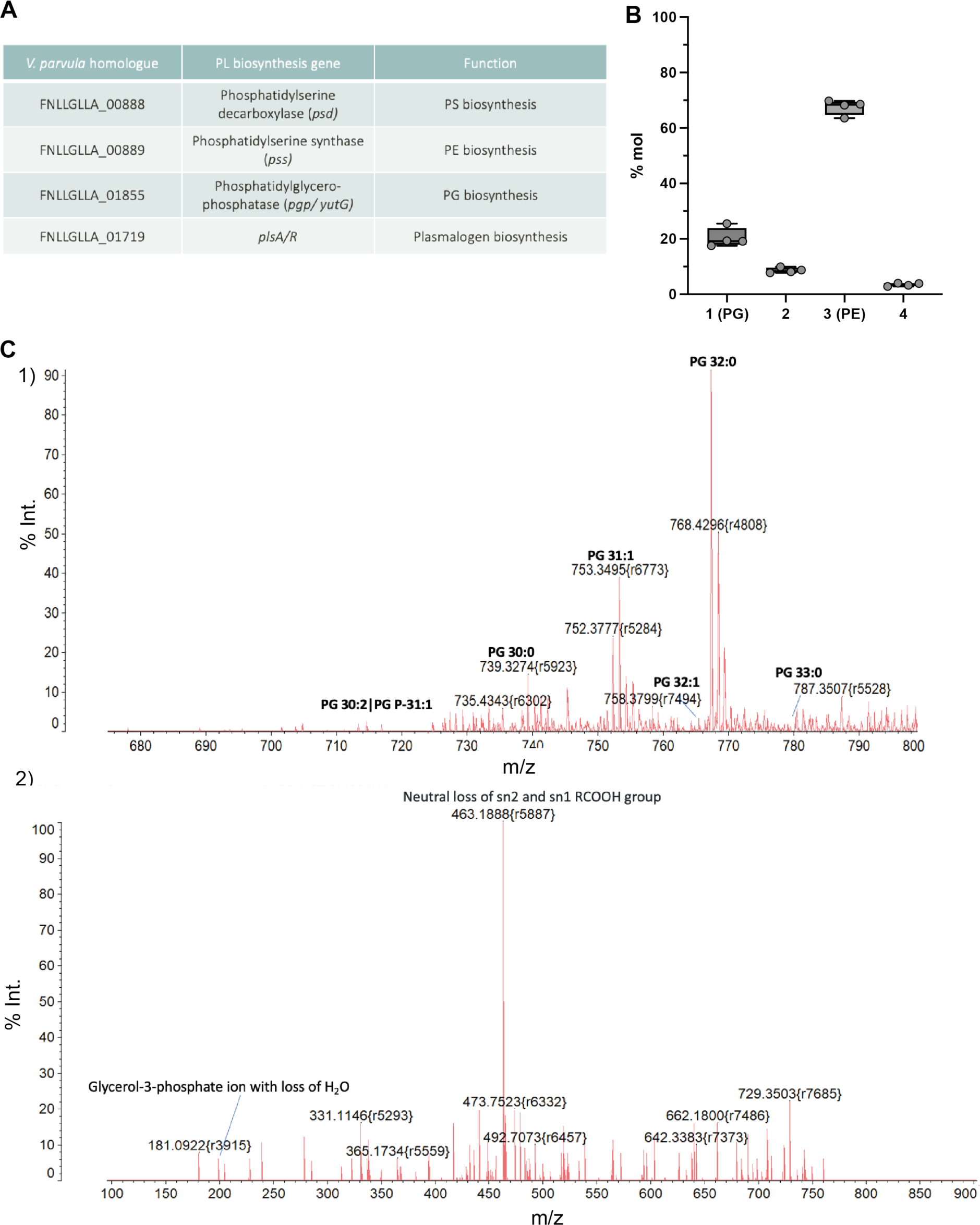
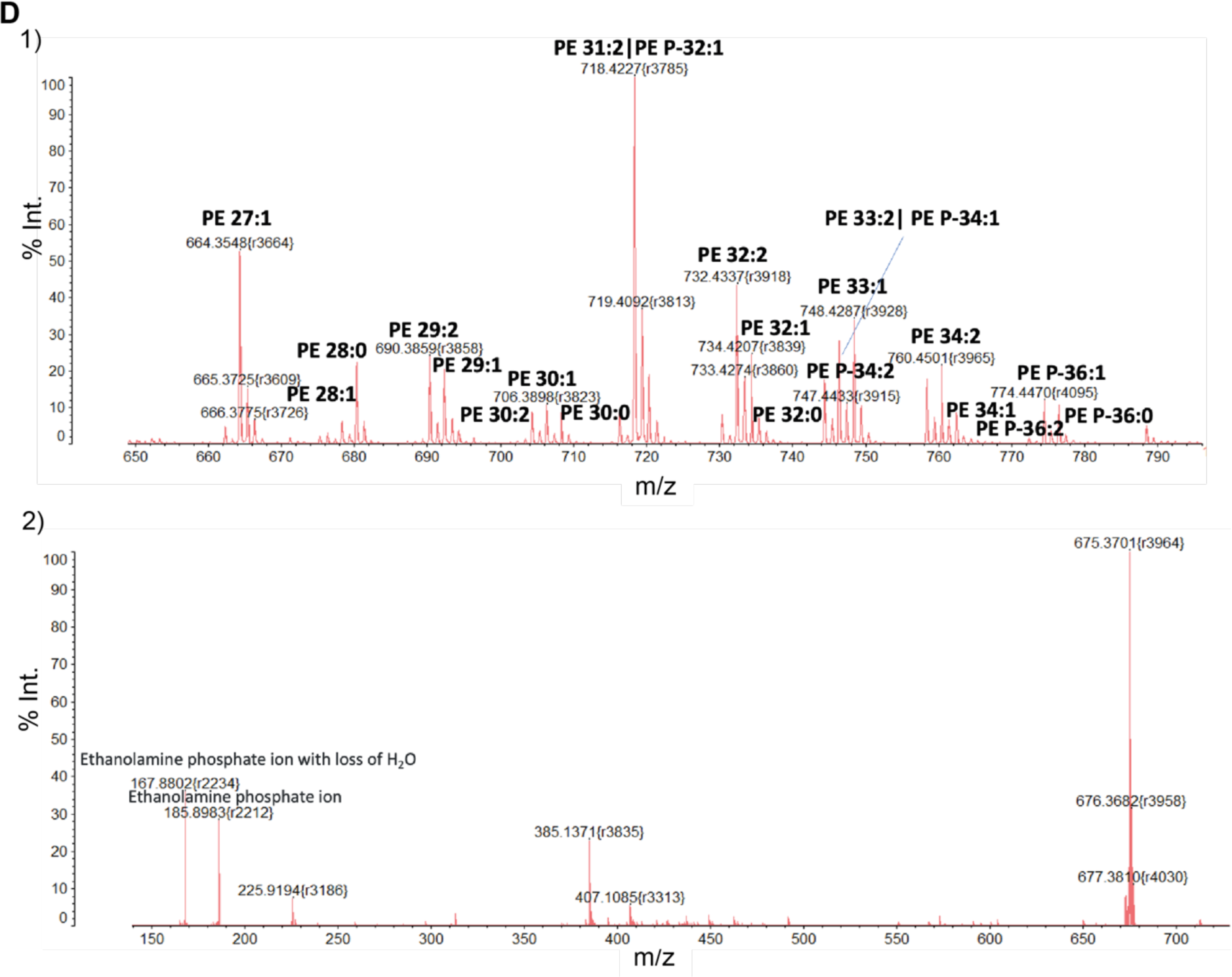
GPL biosynthesis homologues in *V. parvula* and identification of PE and PG. **A) Identification of GPL biosynthesis homologues *in V. parvula***. Homologues of four GPL biosynthesis genes were identified in *V. parvula*, including a homologue for the recently identified operon involved in plasmalogen biosynthesis, *plsAR*. **B) Relative proportion of each lipid species in WT SKV38.** 4 biological replicates were tested, stained with phosphomolybdic acid and quantified by ImageJ; n = 4. **C) Mass spectrometry data – PG identification (band 1).** 1) MALDI-QIT-TOF MS analyses of lipid band 1 (positive ion mode) are shown to illustrate phosphatidylglycerol (PG) diversity from lipid extracts of WT *V. parvula* SKV38. 2) MS/MS spectrum of the main species of PG, PG 32:0 at *m/z* 767.408 with diagnostic ions for PG of 198.9 corresponding to the Glycerol-3-phosphate ion with loss of H_2_O. Only one fatty acid neutral loss was found suggesting the presence of two C16:0. **D) Mass spectrometry data – PE identification (band 3).** 1) MALDI-QIT-TOF MS analyses of lipid band 3 (positive ion mode) are shown to illustrate phosphatidylethanolamine (PE) diversity from lipid extracts of WT *V. parvula* SKV38. 2) MS/MS spectrum of the main species observed by MS, PE 31:2 at *m/z* 718.4 with diagnostic ions for PE of 43 (neutral loss from the precursor ion) and 185 and 167 for respectively the ethanolamine phosphate ion and the ethanolamine phosphate ion with loss of H_2_O.

**Supplementary Table S1.** Lipid assignments of the total lipid extract of *Veillonella parvula*. Analyses by MALDI-QIT-TOF Shimadzu AXIMA Resonance mass spectrometer in the positive mode. The adduct type for phosphoethanolamine (PE) and phosphatidylglycerol (PG) are [M+2Na-H]+.

| Lipids | *m/z* | Assignment |
| --- | --- | --- |
| Phosphatidylglycerol (PG) | 735.434 | PG 30:2 \| PG P-31:1 |
|  | 739.449 | PG 30:0 |
|  | 753.349 | PG 31:0 |
|  | 765.451 | PG 32:1 |
|  | 767.408 | PG 32:0 |
|  | 781.406 | PG 33:0 |
| Phosphatidylethanolamine (PE) | 664.354 | PE 27:1 |
|  | 676.368 | PE 28:2 |
|  | 678.366 | PE 28:1 |
|  | 681.369 | PE 28:0 |
|  | 690.385 | PE 29:2 |
|  | 692.384 | PE 29:1 |
|  | 704.396 | PE 30:2 |
|  | 706.389 | PE 30:1 |
|  | 708.387 | PE 30:0 |
|  | 718.422 | PE 31:2\|PE P-32:1 |
|  | 732.433 | PE 32:2 |
|  | 734.420 | PE 32:1 |
|  | 736.425 | PE 32:0 |
|  | 744.423 | PE P-34:2 |
|  | 746.440 | PE 33:2\| PE P-34:1 |
|  | 748.428 | PE 33:1 |
|  | 760.450 | PE 34:2 |
|  | 762.442 | PE 34:1 |
|  | 772.470 | PE P-36-2 |
|  | 774.447 | PE P-36:1 |
|  | 776.457 | PE P-36:0 |

**Supplementary Figure S2:**
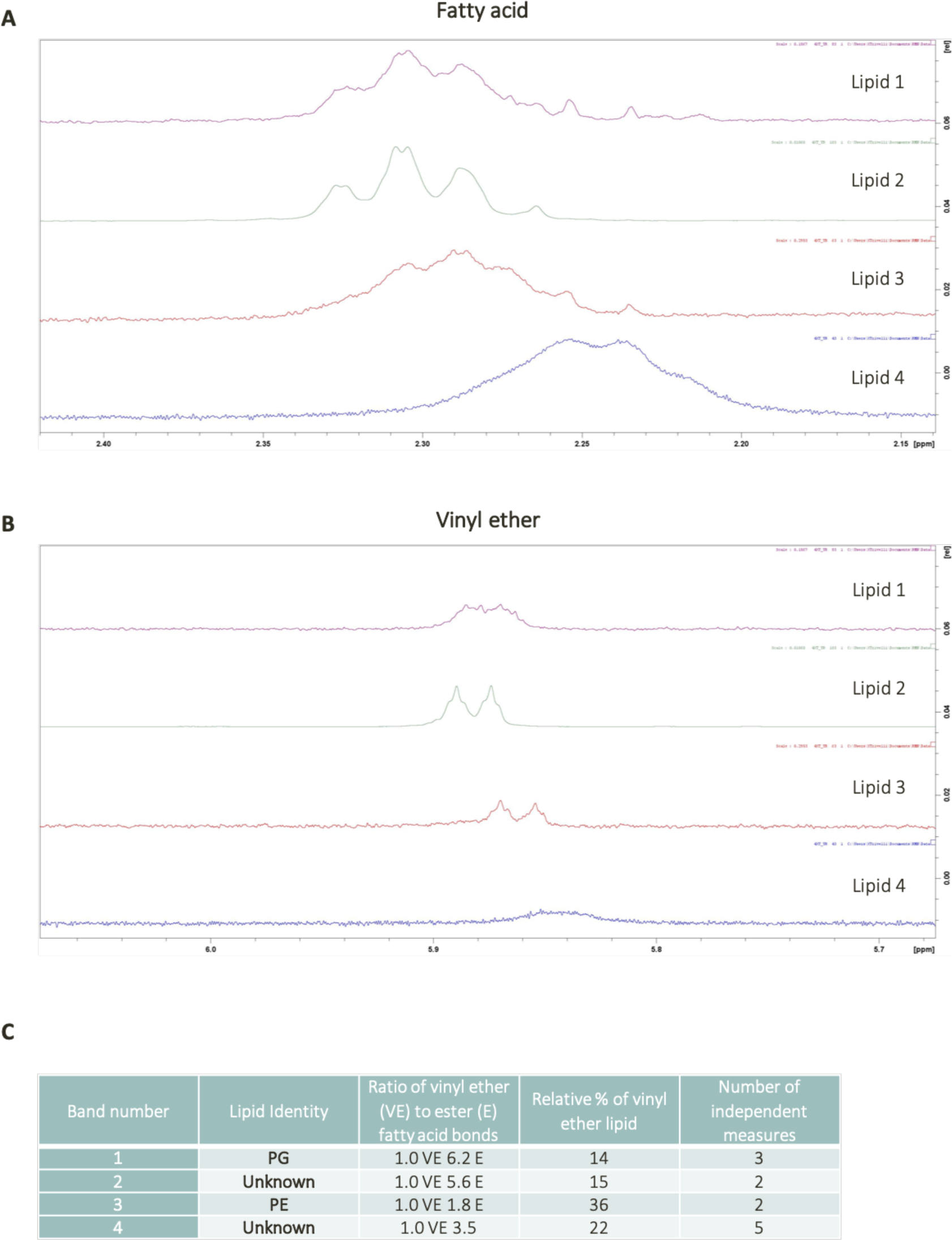
Identification of vinyl ether bonds in *V. parvula* phospholipids. **A)** Proton Nuclear Magnetic Resonance (^1^H-NMR) spectra of lipid fractions 1-4 used for the relative quantification of Ester moieties (FA). Relative quantification was made using signal area from the alpha-methylene of the fatty acids (FA) (two ^1^H signals at ca. 2.30 ppm per FA). **B)** ^1^H-NMR spectra for relative quantification of Vinyl Ether (VE) bonds. Relative quantification was made using signal area for the VE moiety (one ^1^H at 5.88 ppm per VE chain). **C)** Relative proportion of plasmalogen form of each lipid. Based on the relative molar quantity of each of the four major lipids present in the WT, and the NMR data, we calculated the relative molar proportion of lipids containing ether bonds.

**Supplementary Figure S3:**
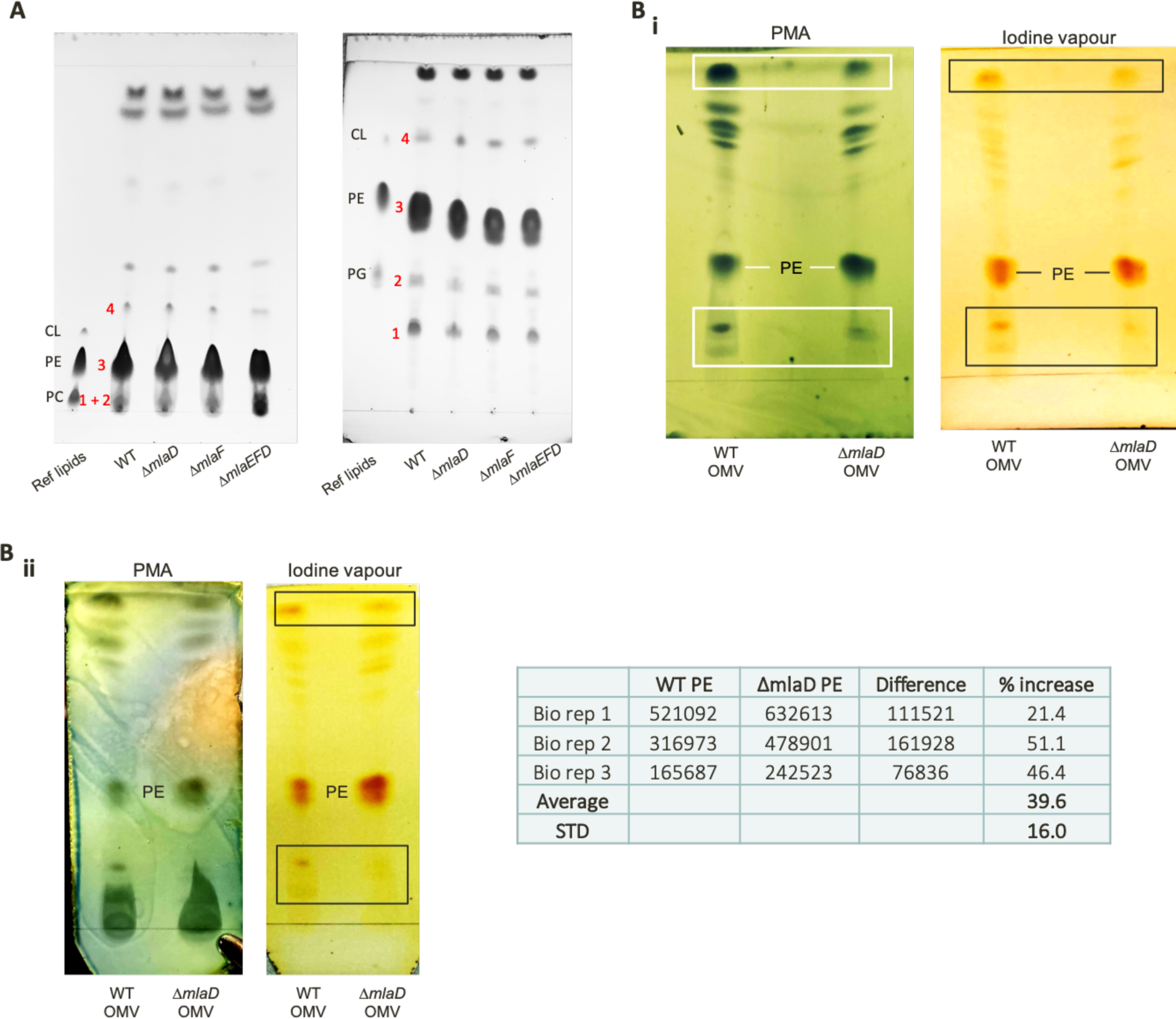
Phospholipid compositions of WT and Δ*mla V. parvula* mutants. **A) Thin Layer Chromatography (TLC) of extracted lipids with iodine vapour staining. i)** Solvent system was chloroform/methanol/water (v/v/v 80:20:2.5). A strong clustering of polar species was observed near the origin of the plate. **ii)** Solvent system was chloroform/methanol/water (v/v/v 65:25:4). This solvent system enabled a better separation of the more polar species, allowing the separation of PE as a double band, likely representative of two different PE species. The major lipid species in *V. parvula*, listed in the table below, are numbered 1 – 4. Commercial reference lipid standards (PC, PE and CL) are labelled. **B) TLC of lipids extracts from OMVs: Phosphomolybdic acid (PMA) and iodine vapour staining (**chloroform/methanol/water 80:20:2.5 v/v/v). Lipid extraction was performed from purified OMVs of WT and Δ*mlaD* strains in biological triplicate as described in the methods. 10μl of extracted lipids were run on TLC, revealing a ∼40% (±16%) enrichment of PE in all Δ*mlaD* OMV lipid extracts. PMA staining of lipid extracts from WT and Δ*mla* OMVs shows a decrease in the relative quantities of two unknown lipid species, highlighted in white. This decrease was also visualised after iodine vapour staining, as highlighted in black. The relative decrease in these two species from the Δ*mla* OMVs confirmed the relative enrichment of PE. **B ii)** Additional biological replicate of OMV lipid extraction and staining, including a table showing ImageJ quantifications of staining intensity of iodine vapour-stained lipid extracts across 3 biological replicates.

**Supplementary Figure S4:**
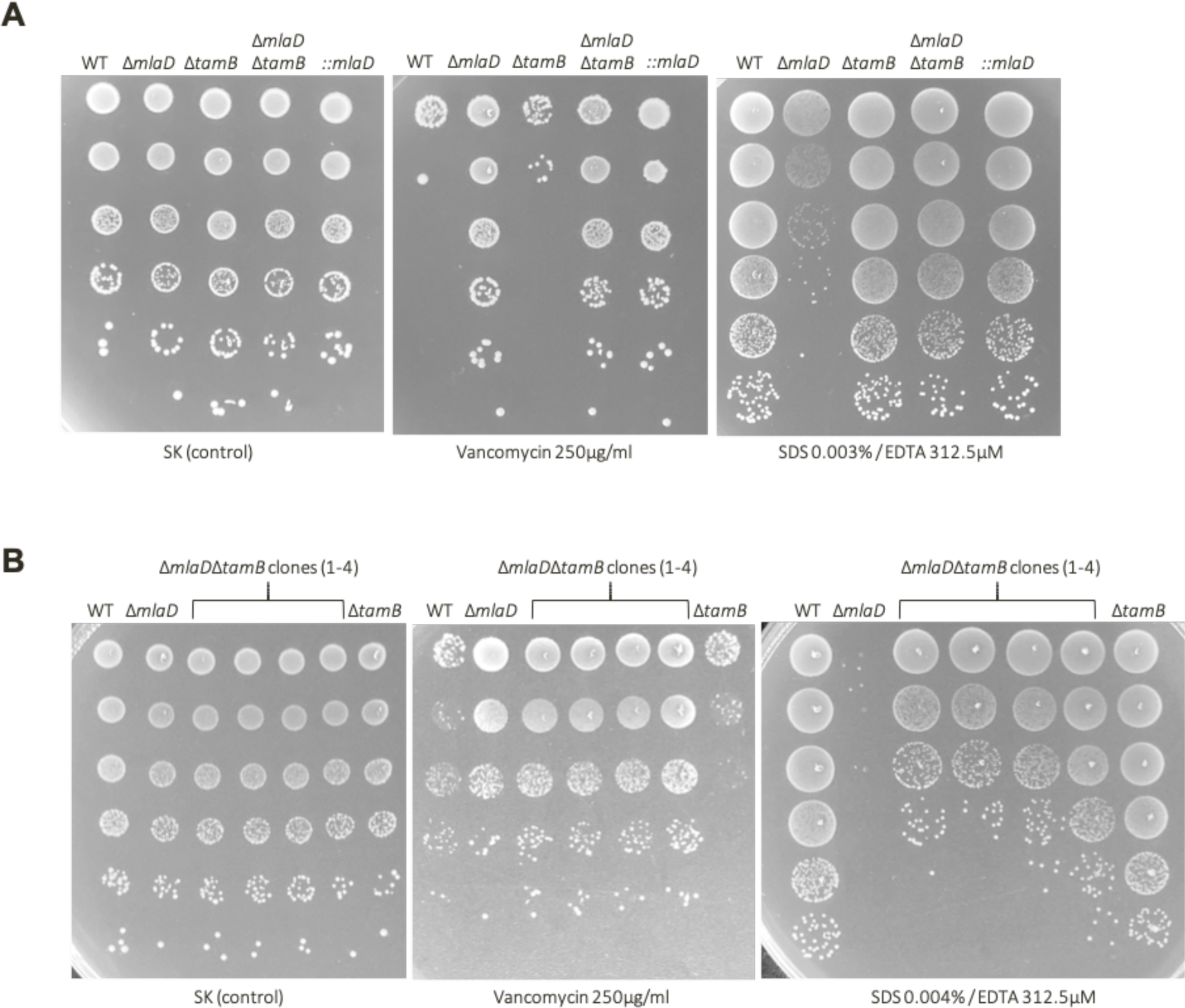
OM permeability phenotypes of Δ*mlaD, ΔtamB* and Δ*mlaDΔtamB* strains. **A)** Overnight cultures of WT, Δ*mlaD, ΔtamB,* Δ*mlaDΔtamB* and Δ*mlaD::mlaD* were plated onto SDS / EDTA and vancomycin to assess OM permeability. At this concentration of SDS (0.003%) and EDTA (312.5µM), the rescue of detergent hypersensitivity of Δ*mlaD* by deletion of *tamB* is striking, and similar to the rescue obtained from reintroducing the expression of *mlaD* itself. Δ*tamB* strains are also highly sensitive to vancomycin. **B)** Cultures of WT, Δ*mlaD, ΔtamB* and 4 independent biological replicates of the double Δ*mlaDΔtamB* mutant were plated onto SDS 0.004% / EDTA 312.5µM and vancomycin 250µg/ml. The slight increase in vancomycin sensitivity of the Δ*tamB* can be observed as compared to the WT, which is in direct contrast to the high resistance of Δ*mlaD* to vancomycin.

**Supplementary Figure S5:**
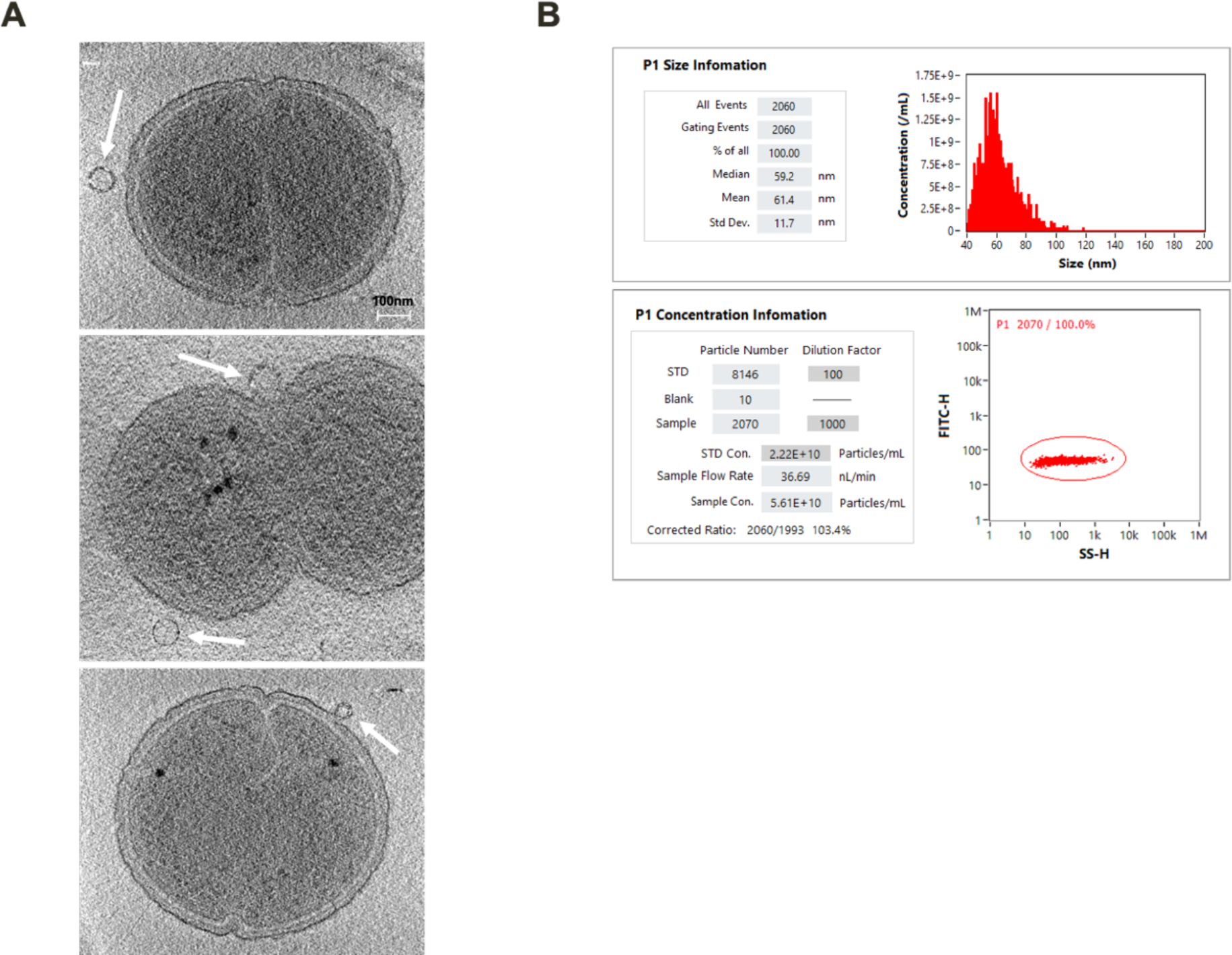
Average size of outer membrane vesicles (OMVs) is consistent across whole cell cultures and purified OMV samples. **A) Cryo-electron tomography images of Δ*mlaD* with outer membrane vesicles (OMVs)**. Cultures of WT and Δ*mlaD* strains were observed via cryo-electron tomography (cryo-ET). In all tomograms of Δ*mlaD*, outer membrane vesicles (OMVs) were observed either attached to the cell or externally (highlighted by white arrows). These vesicles were on average ∼60nm in diameter, matching the average size observed of these vesicles (present both in whole-cell cultures and in purified samples) via NanoFCM. Scale bar = 100nm. **B) NanoFCM report showing size distribution of purified OMV sample.** OMVs were purified from large-scale supernatants of WT and Δ*mlaD* cultures. Prior to concentration of these vesicles via ultracentrifugation, a small sample was extracted and processed via NanoFCM, confirming the average size of OMVs is ∼60nm, matching the average size of OMVs present in full cell cultures of WT and Δ*mlaD* strains, and also the average size observed via cryo-ET.

**Supplementary Table S2:**
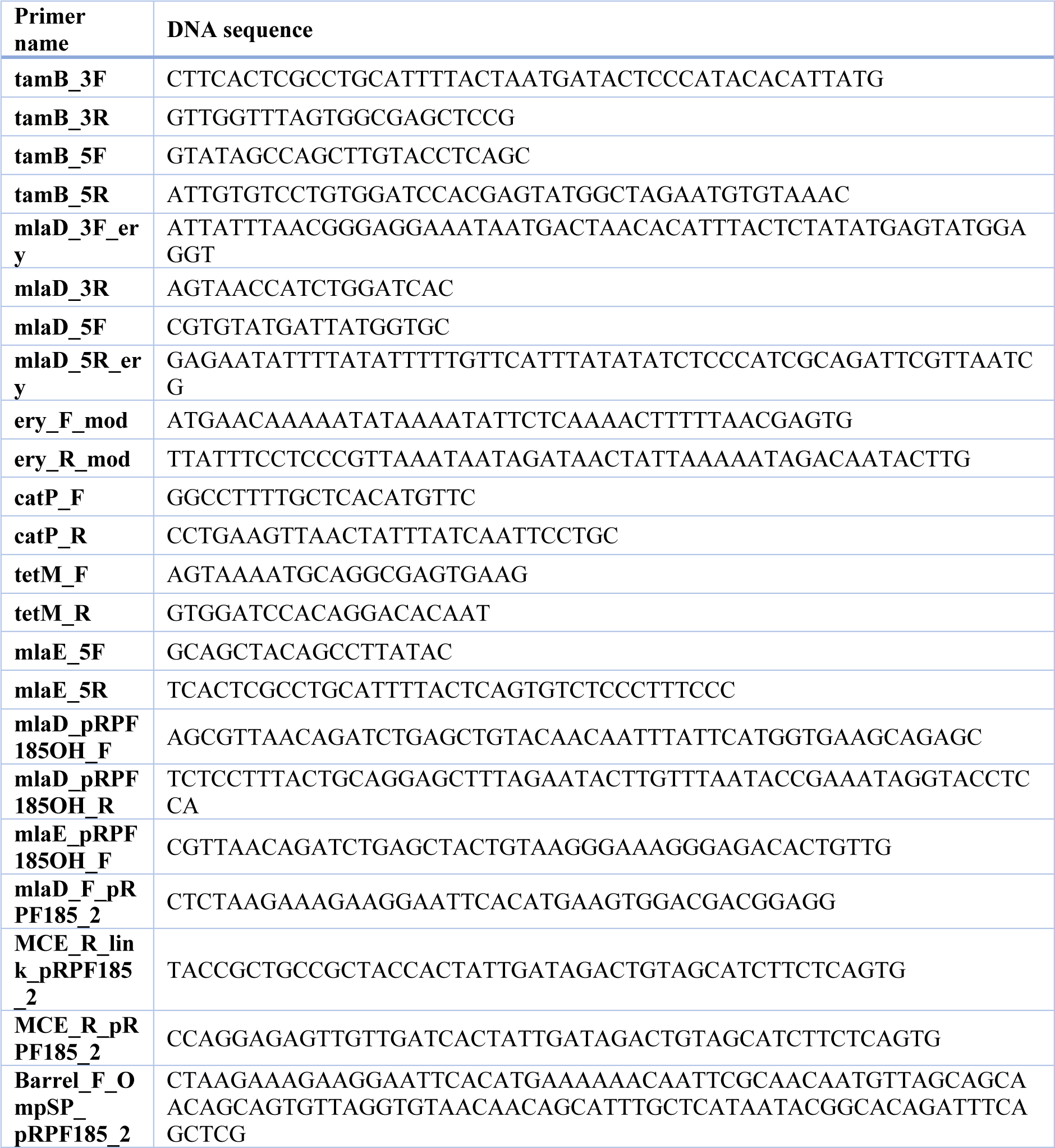
Primers used in this study.

**Supplementary Table S3:** Strains and plasmids used in this study.

| Strain or Plasmid | Description | Antibiotic Resistance | Origin |
| --- | --- | --- | --- |
| SKV38 | Wild-type | / | <sup>1</sup> |
| SKV38$\Delta mlaD$ | Wild-type with chromosomal *mlaD* deletion | Ery | This study |
| SKV38$\Delta mlaF$ | Wild-type with chromosomal *mlaF* deletion | Tc | This study |
| SKV38$\Delta mlaEFD$ | Wild-type with chromosomal *mlaEFD* deletion | Tc | This study |
| SKV38$\Delta mlaD::mlaD$ | $\Delta mlaD$ complemented with pRPF185::*mlaD* | Ery, Cm | This study |
| SKV38$\Delta mlaEFD::mlaEFD$ | $\Delta mlaEFD$ complemented with pRPF185::*mlaEFD* | Tc, Cm | This study |
| SKV38$\Delta mlaD::mlaD$-Barrel-HA | $\Delta mlaD$ complemented with pRPF185::*mlaD* (Barrel: residues 250-419 with C-ter HA-tag with linker) | Ery, Cm | This study |
| SKV38$\Delta mlaD::mlaD$-MCE-HA | $\Delta mlaD$ complemented with pRPF185::*mlaD* (TM + MCE domain: residues 1-155 with C-ter HA-tag with linker) | Ery, Cm | This study |
| SKV38$\Delta tamB$ | Wild-type with chromosomal *tamB* deletion | Tc | This study |
| SKV38$\Delta mlaD\Delta tamB$ | Wild-type with chromosomal *mlaD* and *tamB* deletion | Ery, Tc | This study |
| SKV38$\Delta mlaD$ (Tn) | Wild-type with transposon insertion in *mlaD* homologue | Ery | This study |
| $\Delta mlaD\Delta tamB$ (Tn 27C1) | $\Delta mlaD$ with transposon insertion in *tamB* | Ery, Tc | This study |
| $\Delta mlaD\Delta tamB$ (Tn 90E10) | $\Delta mlaD$ with transposon insertion in *tamB* | Ery, Tc | This study |
| $\Delta mlaD\Delta sstT$ (Tn 5B5) | $\Delta mlaD$ with transposon insertion in *sstT* | Ery, Tc | This study |
| $\Delta mlaD::inter$ (Tn 5A4) | $\Delta mlaD$ with transposon insertion in intergenic region between *sodA* and FNLLGLLA_01854 | Ery, Tc | This study |
| **Plasmids** |  |  |  |
| pRPF215 | mariner Tn delivery plasmid, P<sub>tet</sub>::*HimarI* ITR-*ermB*-ITR *catP* *tetR* | Cm | <sup>2</sup> |
| pRPF185 | Tetracycline-inducible expression system fused with β-glucuronidase *gusA* Term(fdx)-P<sub>tet</sub>-*gusA*-Term(slpA), *catP* | Cm | <sup>3</sup> |
| pRPF185$\Delta gusA_2$ | Tetracycline-inducible expression system with β-glucuronidase *gusA* removed, *catP* | Cm | This study |

